# ProteoBench: the community-curated platform for comparing proteomics data analysis workflows

**DOI:** 10.64898/2025.12.09.692895

**Authors:** Robbe Devreese, Caroline Jachmann, Bart Van Puyvelde, Holda A. Anagho-Mattanovich, Witold E. Wolski, Henry Webel, Matthias Anagho-Mattanovich, Wout Bittremieux, Karima Chaoui, Cristina Chiva, Tine Claeys, Hector Mauricio Castaneda Cortes, Simon Devos, Maarten Dhaenens, Nadezhda T. Doncheva, Viktoria Dorfer, Martin Eisenacher, Ralf Gabriels, Quentin Giai Gianetto, David M. Hollenstein, Lars Juhl Jensen, Vedran Kasalica, Olivier Langella, Caroline Lennartsson, Dominik Lux, Lennart Martens, Mariette Matondo, Teresa Mendes Maia, Emmanuelle Mouton-Barbosa, Alireza Nameni, Michael Lund Nielsen, Jesper Velgaard Olsen, Magnus Palmblad, Christian Panse, Yasset Perez-Riverol, Marina Pominova, Martin Rykær, Eduard Sabidó, Julia Schessner, Martin Schneider, Veit Schwämmle, An Staes, Maximilian T. Strauss, Tim Van Den Bossche, Sam Van Puyenbroeck, Kevin Velghe, Runxuan Zhang, Julian Uszkoreit, Robbin Bouwmeester, Marie Locard-Paulet

## Abstract

Mass spectrometry (MS)-based proteomics is a well-established strategy for analyzing complex biological mixtures. Many MS instruments and data acquisition strategies are available, and the data they acquire differ substantially, thus requiring tailored analysis algorithms. Hence, many dedicated bioinformatics workflows are developed. These are in constant evolution, and the community lacks a centralized platform for comparing their performance. Here, we propose ProteoBench, a single platform that brings together software developers and software users to provide an ever-evolving comparison of state-of-the-art proteomics data processing tools. ProteoBench is an open-source resource that enables the community to evaluate data analysis workflows, develop benchmarking modules dedicated to specific comparisons, and discuss the best methods to compare software tools. The platform ensures that the benchmark evolves alongside advances in proteomics data analysis workflows. ProteoBench guides researchers towards the best-suited tool and parameters for their specific project and data according to their needs, and developers can test their newly developed tools or workflows privately, before adding them as public references. This community-driven effort will increase transparency and reproducibility between MS data analysis workflows, as well as facilitate the development and publication of software workflows in the field.

## MAIN

Mass spectrometry (MS)-based proteomics is an invaluable method for analyzing complex biological mixtures. It is routinely used to compare relative protein quantities in bulk cell lysates, analyze post-translational modifications, identify protein–protein interactions, disease-specific protein variants, and the function and dysfunction of biological processes in general^1^. This variety of applications is accompanied by an increasing number of instruments and data acquisition strategies that generate specific types of data, which in turn require tailored analysis algorithms. For these reasons, numerous workflows exist that are dedicated to the analysis of MS-based proteomics data, and new ones are continually being developed. Information about how these new workflows perform is often incomplete and biased^2^, relying mainly on performance results reported by the developers. Furthermore, their performance can differ substantially from previously published workflows or older versions of the same workflow. This lack of continuous and transparent benchmarking makes it challenging to evaluate how new or updated tools compare with each other.

Even when thorough evaluations are performed on several workflows, these typically isolated efforts can still be problematic. For example, new tools are usually compared to existing tools with similar functionality. In such cases, it has been shown that comparisons often favor the tool described in the paper because the authors have a better knowledge of their workflow (compared to the other tools), as well as a better ability to fix bugs and adapt parameters of their tool^2^. Furthermore, they can choose the evaluation metrics or datasets best suited to their tool. Although this “optimistic bias” is well acknowledged in the community, it is very difficult to avoid. Independent research groups have provided resources to facilitate comparison of MS analysis software, such as samples with known composition that act as a gold standard^3–5^. Additionally, several studies have presented independent benchmarks^6–13^, offering valuable insights to the community. However, these benchmarks are rarely comparable, as each software comparison stands alone and can quickly become obsolete. Beyond published research, many researchers routinely evaluate the performance of new workflows internally, but these findings are seldom made publicly available.

More fundamentally, the field lacks standardized evaluation approaches and consensus metrics, making it difficult to align results across different benchmarking efforts. Thus, even valuable independent studies cannot provide a sustainable or directly comparable framework for the community. This challenge is becoming increasingly urgent as advances in mass spectrometry, including data-independent acquisition, ion mobility separation, and single-cell proteomics, generate increasingly more complex and heterogeneous datasets. These developments further emphasize the need for a continuously updated, community-driven benchmarking platform that can evolve in parallel with rapidly evolving technological developments.

Other fields have already addressed these issues by developing continuous benchmarking platforms such as CAMEO^14^ and CASP^15,16^ for protein structure prediction, and more recently the Open Problems Platform^17^ for downstream single cell genomics analysis tasks, such as batch effect removal. However, there are currently no platforms available that provide comprehensive and detailed evaluations tailored to proteomics data analysis workflows.

ProteoBench (proteobench.readthedocs.io) aims to unify benchmarking efforts into a single platform, fostering collaboration between developers and end-users to facilitate an ongoing comparison of state-of-the-art MS data analysis workflows. As a fully transparent resource, it is dedicated to the standardized comparison of proteomics data analysis software. It is structured into independent benchmark modules that are designed to cover specific aspects of proteomics data analysis. Each ProteoBench module defines a set of pre-established metrics that are used to evaluate workflow performance. These metrics are applied consistently across all workflow results, allowing users to compare them in a standardized way. Each module also provides the input files, ensuring that all workflows are run on the same data. Users can download these files, analyze them using their preferred workflow, and upload the resulting output back to ProteoBench. This allows users to assess their workflow performance in relation to publicly available workflow results contributed by the community. Users can choose to share their results publicly, which permanently adds them to the module’s main figure. This figure visualizes comparative performance across all public submissions, allowing the dataset to evolve over time and reflect the most current tools and software versions.

Anyone can contribute by making workflow results publicly available, proposing new modules, and/or collaborating with the ProteoBench core team on platform development. ProteoBench is therefore an innovative resource for the proteomics community, providing a unified, flexible and open platform for benchmarking. It enables researchers to base workflow selection on systematic evaluation tailored to project and data needs. At the same time, it supports developers by supplying state-of-the-art datasets and a standardized benchmarking framework for testing and improving their tools.

## RESULTS

### Description of the ProteoBench platform

ProteoBench is an open platform for comparing proteomics data analysis workflow performances and a forum for the proteomics community to discuss current benchmarking methods and propose new ones. It contains several benchmark modules, each dedicated to a specific type or step(s) of the data analysis. ProteoBench is accessible via its web application (proteobench.cubimed.rub.de) and a dedicated documentation portal (proteobench.readthedocs.org). Community engagement is coordinated through the ProteoBench GitHub Discussions forum (github.com/orgs/ProteoBench/discussions). There, users can review and debate modules and results, report issues, and propose new modules or platform-wide changes to methods, implementation, and documentation. This open process underscores ProteoBench’s commitment to transparency, collaboration, and community-driven development.

Each benchmarking module is structured as shown in **Figure 1a**. At any time, it is possible to visualize the main module figure that compares the performance of publicly available workflows through the public web interface (“public data visualization”) (**Supplementary Fig 1-2**). Limiting a module’s main figure to only a few metrics makes it possible to present many workflow results in a single summary plot, compactly showing their relative performance. All these results are generated with the data provided in the module (“input data”). These can be downloaded to run any new workflow, and the results can then be uploaded to the ProteoBench module via its web interface (**Supplementary Fig 3**). Results are parsed to produce an intermediate table that presents a given workflow output in a homogenized fashion (same table format for all workflows), with additional module-specific metrics (**Supplementary Fig 4**). These intermediate tables are downloadable and can be easily compared between all workflow outputs. Once benchmark metrics are calculated for this new workflow output, they are added to the main module plot for comparison to other public workflow results (**Supplementary Fig 5**). Furthermore, each workflow output can be explored in depth in a dedicated tab where specific plots allow a thorough inspection of the data (**Supplementary Fig 4**). A pMultiQC^18^ report can be generated for each output and downloaded by the user; providing quality-control metrics and plots, including intensity distributions per run and the number of missing values, among other metrics (an example report can be seen online: pmultiqc.quantms.org/ProteoBench/multiqc_report.html)

**Figure 1:**
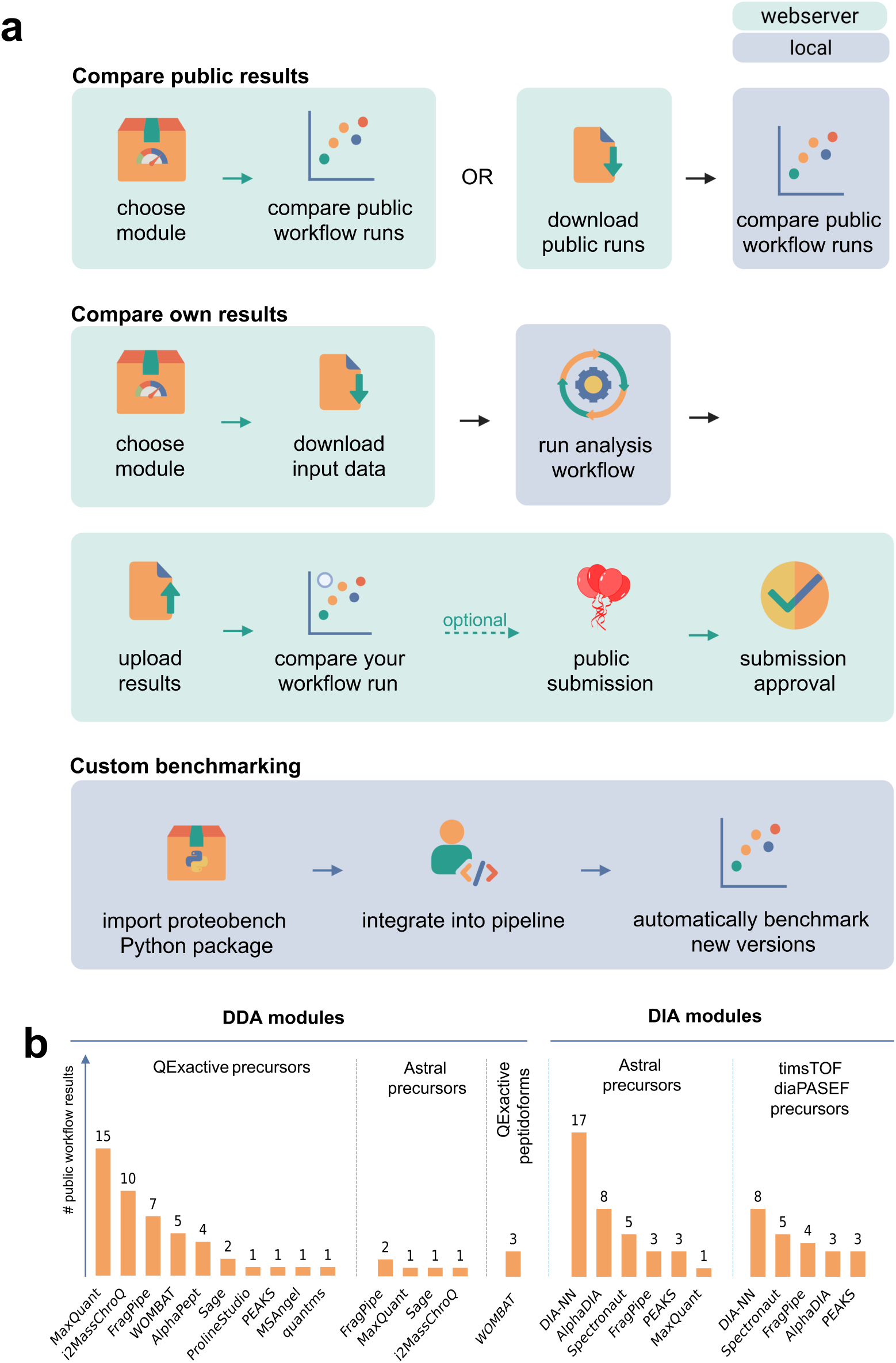
ProteoBench Overview. **a**, The three primary ways to use and interact with ProteoBench. Users can compare workflows using publicly shared results either on the web interface or by downloading them locally. They can also benchmark their own results and, optionally, submit them for publication on the webserver. For reproducibility, public submissions require uploading the parameter file(s) in addition to the results. Workflow developers can integrate the ProteoBench Python package into their projects to streamline benchmarking. Box colors indicate which steps take place on the webserver (green) or locally (blue). **b**, Number of workflow results made public in ProteoBench modules.

After inspecting their uploaded workflow output, users can choose to make their data publicly available by providing the associated parameter file, ensuring full transparency (**Supplementary Fig 6**). The input data, intermediate table, calculated metrics, and parameter files are then inspected by a member of the ProteoBench team using a set of established guidelines (proteobench.readthedocs.io/en/latest/developer-guide/reviewing-new-point-pr/) before being added to the set of publicly available workflow run outputs. Once added to the module’s main figure, these data (raw workflow outputs, associated parameter files, and intermediate metrics calculated by ProteoBench) are fully accessible through the module interface for deeper inspection (**Supplementary Figure 2**).

At the time of writing, ProteoBench has five active modules (**Figure 1b**) and six proposed modules are at various stages of development (**Supplementary Table 1**). In this manuscript, we focus on the active modules. These are dedicated to estimating the depth and error of precursor ion quantification of data-dependent and data-independent acquisition (DDA and DIA, respectively) data acquired with Q Exactive HF-X, Orbitrap Astral, and timsTOF SCP instruments. These modules already present the results of 115 workflow runs from 13 software tools (**Figure 1b**), and indicate differences between workflows and workflow parameters, as discussed in the next paragraphs.

In addition to its public web interface, ProteoBench is also distributed as a PyPI package, making it straightforward to install and integrate into existing Python-based environments. All benchmarking modules and their functionalities are available through this package, including the parsing of output files into standardized intermediate tables, metric calculation, and result visualization. This flexibility enables software developers to incorporate ProteoBench into automated testing environments and continuous integration frameworks, and facilitates more extensive benchmarking analyses. Tutorials on how to use the Python package are available on GitHub (github.com/Proteobench/ProteoBench) in the form of Jupyter Notebooks. Additionally, the ProteoBench web interface can be deployed locally, offering users full agency over their data and computational environment while benefiting from the interactive, browser-based user experience.

### Module for DDA quantification of precursor ions with a mixed-species sample

DDA is a well-established method for global proteomics analysis. Many tools exist to match MS/MS spectra to peptidoforms and determine their relative quantities across MS runs. These tools utilize various algorithms and scoring schemes for MS/MS identification, feature detection, retention time alignment, false discovery rate control, and many other processing steps. As a result, different tools yield varying sets of identified precursor ions and produce different quantification values. To systematically compare workflows for label-free quantification (LFQ) data, we developed a DDA LFQ ion quantification module that evaluates the sensitivity and quantification accuracy for data acquired in DDA on a Q Exactive HF-X Orbitrap (Thermo Fisher Scientific). This module uses six raw files originally presented in a comprehensive benchmark dataset for LFQ analysis^3^. The spectra were acquired on a Thermo Orbitrap QE HF-X coupled to an UltiMate 3000 LC-system, using a 120-minute gradient, fragmentation of the top 12 most intense precursors, and dynamic exclusion within 30 seconds.

The raw files provided with this module represent two experimental conditions (“condition A” and “condition B”) in triplicate and consist of a mixture of tryptic peptides from three species (*Homo sapiens, Saccharomyces cerevisiae,* and *Escherichia coli*). The composition of the peptide mix differs between conditions A and B, with known peptide quantities from each species. Sample A was generated by combining protein digests derived from Human, Yeast, and *E. coli* in proportions of 65%, 30%, and 5% weight-for-weight (w/w), respectively. Sample B was prepared by mixing the same sources at relative abundances of 65% Human, 15% Yeast, and 20% *E. coli* (w/w). These mixtures yield log2-transformed fold changes (log2FC) of 0, −1, and 2 for Human, Yeast, and *E. coli*, respectively.

This experimental design allows for the following metrics to be calculated from workflow results, including only ions that uniquely match a single species:

- Depth(k): Number of precursor ions quantified in at least k runs, where a precursor ion is defined by its sequence, localized modifications, and charge.
- QuantError: Difference between the observed and expected log2FC of a precursor ion. The expected log2FC is determined by the experimental setup, whereas the observed log2FC is calculated by subtracting the mean log2-transformed precursor intensity in condition A from that in condition B. QuantError is denoted as ‘epsilon’ in intermediate files.
- AbsQuantError: Absolute QuantError.
- MedianQuantError(k): Median QuantError of all precursor ions quantified in at least k runs.
- MeanQuantError(k): Mean QuantError of all precursor ions quantified in at least k runs.

The module provides a FASTA file containing sequences of all UniProtKB/Swiss-Prot^19^ proteins from the three species, and the universal protein contaminant database^20^. This ensures that all analyses in this module use the same raw data and FASTA database. Users can download the raw files and FASTA file, perform their searches and quantification using the tools and parameters of their choice, and then upload the results to ProteoBench for comparison. At the time of writing, output files from MaxQuant^21^, FragPipe^22^, ProlineStudio^23^, MSAngel (profiproteomics.fr/ms-angel), OpenMS^24^, WOMBAT-P^25^, AlphaPept^26^, PEAKS Studio^27^, Sage^28^, quantms^29^ and i2MassChroQ^30^ are directly compatible with this module. Additionally, it is possible to use a minimal tab-delimited format including the precursor ion identities (sequence, charge, and localized modifications) and quantities for each run to visualize results from not yet compatible software tools. More information about output format compatibility and workflow-specific instructions can be found in the ProteoBench documentation (proteobench.readthedocs.io).

Based on the user-provided data, ProteoBench calculates the quantification depth (Depth(k)) and global quantification error (MedianQuantError(k) or MeanQuantError(k)) as described above. These metrics are then shown in a single overview plot (**Figure 2a**) that enables direct visual comparison of workflow runs from different tools, between different sets of search and quantification parameters (**Figure 2b**), and between different versions of the same workflow (**Figure 2c**). In the overview plot, users can select either the mean or the median quantification error for the horizontal axis. The mean is more sensitive to outliers, making it a suitable metric when ensuring accurate quantification across all data points is a priority (**Supplementary Figure 7**). In contrast, the median provides a more robust representation of the general trend, as it is less influenced by extreme values (**Supplementary Figure 8**). Users can interactively adjust the filtering threshold using a slider above the plot, which sets the minimum number of MS runs in which an ion must be quantified to be included in the metric calculation (k, default: 3). Stricter filtering, requiring precursor ions to be quantified in more MS runs, reduces the total number of quantified precursor ions (**Supplementary Figures 7-8**).

**Figure 2:**
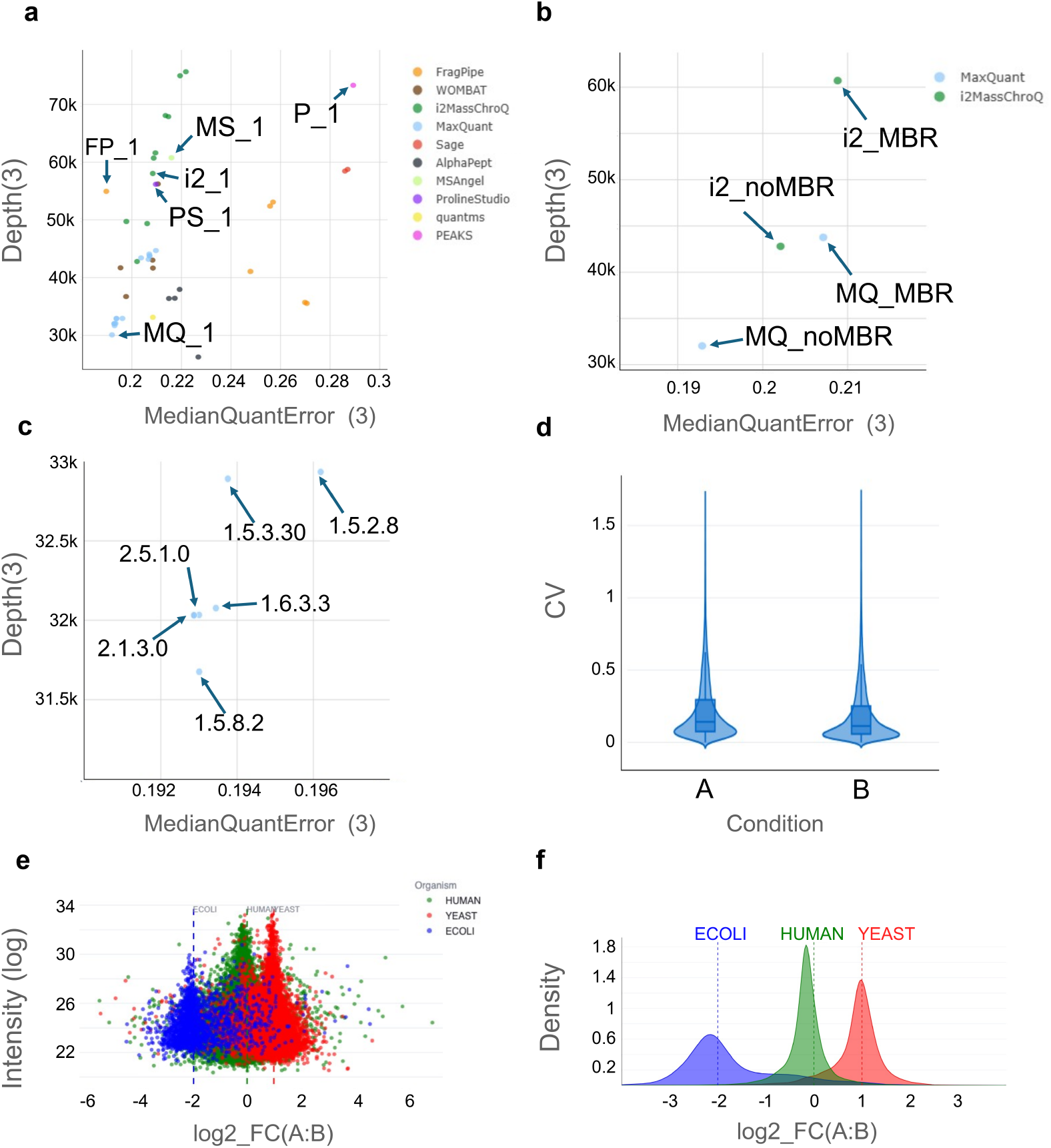
DDA Ion Quantification Benchmarking plots. **a**, Main figure of the DDA ion quantification module: a scatterplot showing median quantification errors of all precursor ions quantified in at least 3 out of 6 MS runs [MedianQuantError(3)] plotted against the number of quantified precursor ions [Depth(k)]. Quantification error is calculated as the median difference between the observed log2FC and the expected log2FC. This plot is interactive and can be filtered by the minimum number of MS runs in which an ion is quantified (k). Additionally, it can be switched to display the mean or median quantification error. **b-c**, Main module plot with a subset of results that illustrate the impact of enabling match between run (MBR) for MaxQuant and i2MassChroQ **(b)** and differences between MaxQuant versions **(c)**. “i2_noMBR” and “i2_MBR” are i2MassChroQ runs without and with MBR, respectively; “MQ_noMBR” and “MQ_MBR” are MaxQuant runs without and with MBR, respectively. Parameters are listed in **Supplementary Table 2**. **d-f**, Example of in-depth plots generated by ProteoBench with the i2MassChroQ workflow result “i2_1”: coefficients of variation within 3 replicates of condition A and B (d); MA plot showing the mean precursor ion log₂-transformed intensities vs. their log2FC (e); Species-specific precursor log2FC distributions (f). In (e) and (f), vertical lines correspond to expected log2FCs between conditions A and B, and Human, Yeast and *E. coli* precursors are presented in green, red and blue, respectively.

**Figure 2b** shows that enabling match between runs (MBR) leads to a 27% increase in precursors quantified in a minimum of 3 runs (Depth(3)) in MaxQuant, which is accompanied by a 7% increase in MedianQuantError(3). With i2MassChroQ, enabling MBR results in a 30% increase in Depth(3) associated with a 3% increase in MedianQuantError(3). This slight increase in global quantification error mainly stems from the precursor ions quantified with MBR that have poor quantification quality (**Supplementary Figures 9-10**). This higher quantification error is the result of an increased risk of incorrectly matching MS1 peaks during the MBR process. Because publicly available workflow outputs include older versions of MaxQuant, we see that using different versions of MaxQuant slightly impacts the results. For example, using the same input files and parameters results in a slightly lower number of quantified precursors in version 1.5.8.2 (Depth(3)=31,676) than in version 1.5.2.8 (Depth(3)=32,935), but the quantification error stays mostly the same (MedianQuantError(3)=0.193 and 0.196, respectively) (**Figure 2c**).

In addition to the overview plot, three other plots are generated to assess the performance of individual workflow results. A violin plot shows the reproducibility of quantification as the coefficient of variation (CV) calculated from the precursor ion intensities across the replicates (excluding contaminants and peptides matched to more than one species, and including precursor ions quantified in 2-3 replicates) (**Figure 2d**). An MA plot shows the relationship between quantification error and average intensity for each precursor ion, grouped by organism (**Figure 2e**). Finally, a density plot of all precursor ions log2FC between condition A and B per species shows the distribution of the observed ratios relative to the expected value (**Figure 2f**).

Beyond visualizing data in the modules’ main plot, ProteoBench provides entry points for users interested in more customized and automated analyses. Users can either start from the intermediate files available for download on the webpage or use the Python package. The Python package allows users to easily parse search engine outputs, calculate metrics (including additional metrics such as median CVs) and visualize them. An illustrative analysis of agreement and disagreement across workflows is shown in **Figure 3**. Of 87,338 precursor ions, 25,334 (29%) were quantified by all workflows. PEAKS and i2MassChroQ contributed the largest and second-largest sets of precursor ions unique to a single workflow, respectively (**Figure 3a**). MaxQuant predominantly quantified ions that were also detected by all other workflows, whereas a substantial number of ions quantified by all other workflows were not captured by MaxQuant. Precursor ions quantified by only one workflow exhibited higher quantification errors than those quantified by all workflows, with the highest errors observed for ions unique to PEAKS (**Figure 3b**). Pairwise comparisons across workflows are provided in **Supplementary Figure 11**.

**Figure 3:**
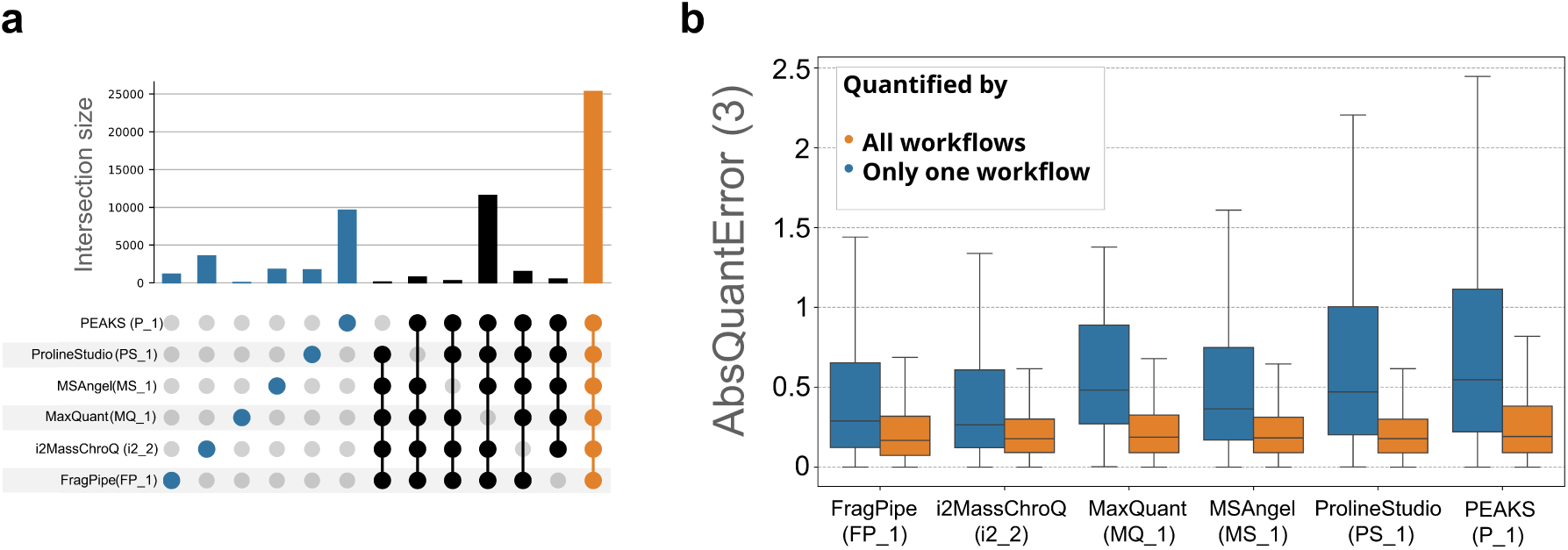
Local analysis of benchmarking results using the ProteoBench Python package. **a**, UpSet plot showing the number of precursor ions quantified with only one workflow (blue), all but one workflow (black), and all workflows (orange). **b**, Boxplots of the AbsQuantErrors per workflow for precursor ions quantified uniquely with one workflow (blue) or in all workflows (orange). For both subfigures, k was set to 3. Exact parameters for the workflow runs are listed in Supplementary Table 2 where the column “article_id” contains the identifier associated with the workflow.

At the time of writing, 46 workflow runs from 10 software tools are public and show a diversity in identification and quantification performances that are obtained with the same set of raw files and using the same sequence database (**Figure 1b**). We expect the number of workflow runs to increase over time, as both users and developers will continuously add new workflow runs based on new releases of existing software, newly developed tools, and alternative parameter settings.

Modules for DIA quantification of precursor ions with a mixed-species sample DIA methods have gained significant popularity over DDA and are now widely used across many proteomics laboratories. However, due to their relatively recent emergence and the variety of DIA strategies in terms of acquisition and data analysis, the software tools dedicated to DIA analysis are varied and challenging to compare. Several independent studies have performed a comparison of DIA analysis tools^9,11,13^, but these comparisons quickly become outdated due to the rapid evolution of DIA software. ProteoBench offers a platform for continuously comparing the latest DIA analysis workflows. We aim to develop several modules to accommodate a wide range of datasets, as it is essential to know whether workflows and specific software parameters perform better or worse depending on the sample type, data acquisition method, and instrument.

To this end, we have already developed two distinct modules analogous to the DDA quantification module as outlined in the previous section. Raw MS data were acquired in DIA mode with the Orbitrap Astral (Thermo Fisher Scientific) and the timsTOF SCP (Bruker). The benchmark strategy, which involves using varying mixtures of different organisms, is identical to the one used for the DDA module^3^, and is described in the Methods section as well as in each module’s documentation. The raw data were acquired for the purpose of these benchmarking modules (see Methods). The calculated metrics and functionality of these modules are equivalent to the aforementioned DDA quantification module. At the time of writing, the DIA modules support output files from AlphaDIA^31^, DIA-NN^32^, FragPipe^33^, MaxQuant (MaxDIA)^34^, PEAKS Studio^27^, and Spectronaut (biognosys.com/software/spectronaut).

As of this writing, the Astral and timsTOF DIA modules each include 30 and 23 public workflow runs, respectively, derived from five different software tools (**Figure 4a** and **b, Supplementary Figure 12 and 13**). Here, we describe an in-depth benchmark of a limited set of public workflow runs that share a standard set of parameters (**Supplementary Table 2**; all parameters not mentioned were left to default values). Here, 44% of all precursors (69,673 out of 156,629 with k=3) were quantified across all workflows runs, with DIA-NN (D_A) and PEAKS Studio (P_A) quantifying the highest number of precursors (125,114 and 125,026 respectively with k=3) in the Astral module (**Figure 4c**). However, in the timsTOF module, the highest number of precursors are quantified by Spectronaut (S_T) (125,604 with k=3), underscoring the value of evaluating performance with datasets originating from different instruments. As in the DDA module, we observe higher quantification errors for precursors quantified by a single workflow compared to those quantified by all workflows (**Figure 4d**).

**Figure 4:**
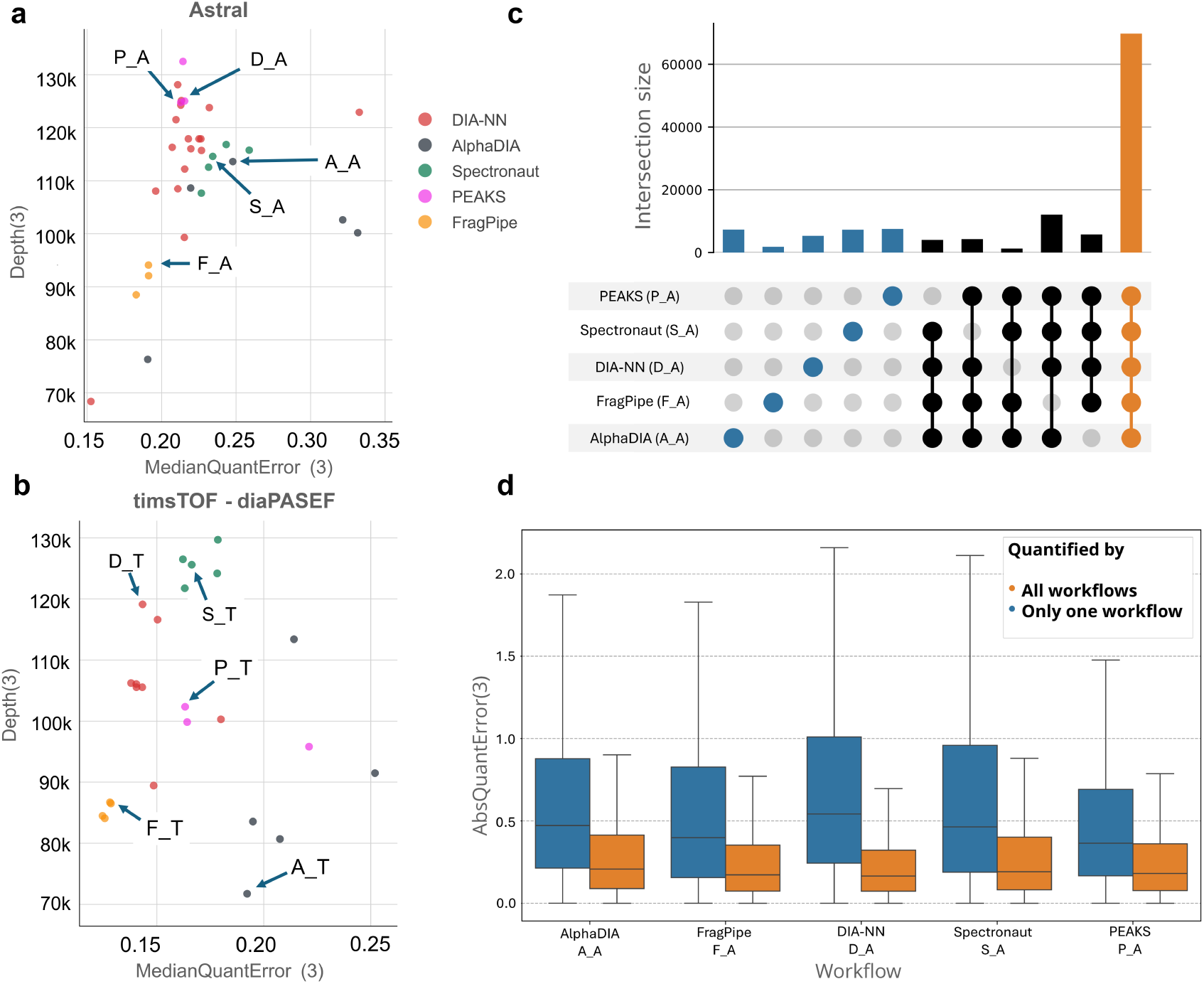
ProteoBench modules for comparison of precursor ion LFQ in DIA data. **a-b**, Main module figure showing the relationship between the number of quantified precursor ions and the median quantification error for the Astral (**a**) and timsTOF SCP (**b**) DIA modules for k = 3, where k is the minimum number of runs in which a precursor ion is quantified. The workflow runs with a standard set of parameters that are annotated with an identifier in Supplementary Table S2 detailing all parameters. **c**, UpSet plot showing the number of precursor ions quantified with only one workflow (blue), all but one workflow (black), and all workflows (orange) in the different DIA workflows compared. **d**, Boxplots of the precursor ions AbsQuantErrors per tool for precursor ions quantified uniquely with one workflow (blue) or in all workflows (orange) in the Orbitrap Astral data.

All tools produced log2FCs centered around the expected ratios, though with noticeably varying levels of distribution spread and skewness (**Supplementary Figure 14**). These discrepancies reveal that tools not only differ in depth but also in the reliability of their quantification. Interestingly, the quantification of PEAKS showed the most narrow distributions around the expected ratios in the Astral module, whereas PEAKS showed the weakest quantitative performance in the DDA module (**Figure 3b**). This further emphasizes how performance can depend on the acquisition strategy. Furthermore, some precursor ions showed measured ratios that deviated from expected values. In some cases, these ions were quantified inconsistently across workflows, indicating workflow-specific challenges in accurately capturing their signals. In other cases, several workflows returned similar values that nevertheless differ from the expected ratios, which may indicate commonalities in algorithmic approaches (for example between DIA-NN and FragPipe, which uses DIA-NN as a quantification engine) or broader, workflow-independent effects. These observations are particularly evident for subsets of *E. coli* precursors and underscore how ProteoBench can be used to identify systematic quantification biases that are either unique to individual tools or shared across multiple tools.

These instrument-specific DIA modules highlight performance differences between workflows across datasets generated by different instruments. By incorporating multiple datasets, the DIA modules in ProteoBench provide a robust framework for evaluating current analysis methods. As DIA acquisition strategies and software for DIA data analysis continue to evolve, we aim to expand ProteoBench with additional modules to provide a broadly applicable, standardized, community-driven benchmarking.

## DISCUSSION

ProteoBench is a community-driven effort to continuously compare proteomics data analysis workflows. It helps users both visualize differences between workflows at a glance and perform in-depth, customizable comparisons using the Python package. It enables the proteomics community to quickly perform tool and parameter comparisons, identify problems with software releases or parameter choice, and gain valuable insights for improving workflows.

In this manuscript, we present how it can be used to easily compare workflows with different software tools, software versions, and parameters. In the discussed modules, we observe a general trade-off between sensitivity and global accuracy, where an increase in the number of quantified precursors comes at the cost of including more poorly-quantified precursor ions.

ProteoBench deliberately shows a single, summarized plot for comparing workflows to prevent information overload. This approach ensures that the main plot of each module provides a clear and easy-to-read reference for the community. While reducing the number of metrics prevents an in-depth evaluation of differences between workflows, the design of ProteoBench allows for further comparison of workflows by making all input, intermediate, and output data from ProteoBench downloadable. To facilitate deeper analysis and visualization of these data, we provide publicly available Jupyter Notebooks (github.com/Proteobench/ProteoBench/tree/main/jupyter_notebooks) that contain workflow-specific visualizations as presented in **Figure 2d-f**.

ProteoBench enables an easy, flexible, and timely comparison of existing data analysis workflows, providing a frame of reference for newly developed workflows that evolves continuously, according to the needs of the field. There are a few things that ProteoBench is not designed for: it does not point to a single best one-size-fits-all data analysis workflow; it should not be used as evidence for generalized statements about a workflow’s performance; and should not be used by developers as a single performance measure of their workflow. Moreover, we are aware of the possibility that developers may overfit a tool to a given module and its dataset. We believe that this is mitigated by two important features of the platform:

1. All workflow runs are associated with all the information necessary to be completely reproduced. All parameters are publicly available, and anyone can reproduce a given workflow or test different parameters, provided that they have access to the software tool.
2. The same workflow can be applied to datasets from different modules to evaluate the impact of the input data on the workflow performance. Indeed, ProteoBench comparisons are not constrained by the current evaluation datasets since the community is encouraged to propose new modules that evaluate workflows on diverse sample types and acquisition methods with a variety of evaluation metrics. It is unlikely that a workflow is simultaneously overfitted on several different modules at the same time.

A few modules contain data points corresponding to workflows already benchmarked in other, more thorough comparisons like some of the WOMBAT-P runs^25^ in the modules dedicated to DDA data analysis. Although less detailed, the comparison performed in ProteoBench includes more workflow outputs, and lasts beyond the date of publication since these points can now be compared to any future runs performed. Other modules are in preparation that will be based on published rigorous benchmarks: one will be dedicated to the comparison of *de novo* search engines^35,36^, the other relies on co-abundance profiles from subcellular fractions of the same sample^37^. Both modules were proposed by people from outside of the ProteoBench core team, which illustrates community involvement.

It is exciting to note that virtually any step of the full proteomics data analysis process can be easily supported in the future. This ease of extending to any analysis workflow is due to the extendable design of ProteoBench, and currently several new modules are being investigated: target-decoy peptide-spectrum match discrimination, protein inference and summarization, post-translational modification localization, or statistical analysis of differentially regulated protein groups, to name only a few. Furthermore, comparisons can go beyond MS-based proteomics and cover other proteomics technologies, such as single molecule sequencing and protein assays. The first ProteoBench modules consist of relatively simple comparisons: sensitivity and global quantification errors are straightforward to calculate. There are numerous workflows that will be harder to compare due to lack of ground truth data, lack of community agreement on the metrics to report, or because workflow outputs cannot be summarized into a reduced number of metrics. ProteoBench provides a place for the community to have these discussions and tackle these challenges together. We believe that the flexibility and open nature of this platform makes it the perfect community tool for both wet-lab scientists and developers to finally compare proteomics software tools, workflows, and analysis steps together.

## METHODS

### ProteoBench workflow

ProteoBench is structured as a public GitHub organization comprising multiple public repositories (https://github.com/Proteobench/). This includes the main ProteoBench codebase that is implemented as a modular Python-based framework. There are additional repositories, each dedicated to manage publicly submitted benchmark results per module.

Each module is tailored to a specific aspect of proteomics data analysis, and has an associated set of downloadable data. Users can interact with the platform through a Streamlit-based web interface, which provides access to benchmarking results and facilitates submission of new workflow outputs.

Upon submission of workflow results, ProteoBench parses the data using a tool- and module-specific settings file. This parsing process converts the input into a standardized, generic format suitable for downstream processing. Once formatted, the parsed data can be compared more easily between workflow tools. These generically formatted results are then used to compute module-specific metrics and generate a corresponding datapoint that summarizes workflow performance, which is subsequently used to visualize comparisons to all publicly available data points.

After submission, the user can choose to contribute this datapoint to the public repository. This submission also requires a metadata file containing all parameters of their workflow. This metadata file can be in the form of a workflow-specific log or settings file. This file is also parsed to extract relevant settings, which are subsequently used to annotate the datapoint. These parameters provide contextual information and are visualized on the public results interface and downloadable in a table format (**Supplementary Table 2**).

After submission, the platform automatically generates a pull request in the corresponding module-specific GitHub repository. This pull request includes a JSON file detailing the new data point, updating the repository’s existing dataset. Following approval by a ProteoBench maintainer, the new datapoint is made publicly accessible via the web interface.

ProteoBench is licensed under the Apache 2.0 license, encouraging contributions from the proteomics community to expand the library of benchmarking workflows, results and datasets.

### Benchmarking procedure of ProteoBench modules discussed in this manuscript

The ProteoBench modules outlined in this article evaluate quantification performance across workflows by processing software tools output files and computing key benchmarking metrics, using a three-species benchmark dataset in both DDA and DIA. These are all designed the same way, and based on the same ground truth. The way ProteoBench calculates metrics, and the associated plots are the same and are described in this section.

First, the precursor-level quantification matrix produced by the workflow is submitted locally to the ProteoBench module, and then parsed using workflow-specific parsers to generate a standardized format. In this format, LC-MS/MS run names are homogenized, decoy peptides and peptides assigned to contaminant proteins are removed, as well as peptides assigned to proteins from multiple species. Precursor ions are then converted to comply with ProForma v2.0 notation^38^. Next, quantification metrics are computed from this standardized format. Precursor ions are grouped by experimental condition, and intensities are log₂ transformed. For each precursor ion in each condition, the mean and standard deviation of both raw and log₂-transformed intensities are calculated, along with the summed intensity within each condition. The coefficient of variation (CV) is determined per precursor ion within each condition if the precursor ion was quantified in at least two replicates per condition, and the log₂ intensity difference between conditions is computed (log2FC). Based on the inferred protein and corresponding species, the difference between the measured and expected log2FC between conditions (QuantError, also referred to as epsilon) is derived for each precursor ion. Metrics are calculated six times, each time for precursor ions quantified in both conditions and a minimum of one (k = 1) to six (k = 6) replicates. These metrics include the median and mean absolute QuantError (referred to as Median- and MeanQuantError(k), respectively), the variance of epsilon, and key CV quantiles (median, 0.75, 0.90, and 0.95).

In the ProteoBench web interface of these modules, data visualization is performed using Plotly (v6.2.0), generating the histograms of log₂ precursor intensity differences between conditions and violin plots of CV distributions. The main benchmarking plot displays the Mean-or MedianQuantError(k) (user-defined) on the horizontal axis against the Depth(k) on the vertical axis, with points colored according to the software tool being benchmarked.

### Data generation for the DIA modules

For the DIA LFQ precursor ion modules, we generated raw files similar to those presented in Van Puyvelde *et al.*^3^ on a timsTOF SCP (Bruker) and Orbitrap Astral (Thermo Fisher Scientific). MS-compatible Human K562 (P/N: V6951) and Yeast (P/N: V7461) protein digest extracts were purchased from Promega (Madison, Wisconsin, United States). The lyophilized MassPrep *Escherichia coli* digest standard (P/N:186003196) was purchased from Waters Corporation (Milford, Massachusetts, United States). All extracts were reduced with dithiothreitol (DTT), alkylated with iodoacetamide (IAA) and digested with sequencing grade Trypsin(-LysC) by the respective manufacturers. The digested protein extracts were reconstituted in a mixture of 0.1% formic acid in water.

For each module, two master samples A and B were created independently, similar to Navarro *et al*.^7^. Sample A was prepared by mixing 65% Human, 30% Yeast, and 5% *E. coli* protein digests (*w*/*w*/*w*). Sample B was prepared by mixing 65% Human, 15% Yeast, and 20% *E. coli* protein digests (*w*/*w*/*w*). When comparing condition A with condition B, the resulting samples have log₂-transformed fold changes (A:B) of 0, −1 and 2 for Human, Yeast, and *E. coli*, respectively.

#### diaPASEF

The resulting peptides were analyzed in triplicate (25 ng) by nanoLC-MS/MS using an UltiMate 3000 RS nanoLC system (Thermo Fisher Scientific) coupled to a timsTOF SCP mass spectrometer (Bruker). Peptides were separated on a C18 Aurora column (25cm x 75 µm ID, IonOpticks) using a gradient ramping from 2% to 20% of B in 30 min, then to 37% of B in 3 min and to 85% of B in 2 min (solvent A: 0.1% formic acid in H2O; solvent B: 0.1% FA in acetonitrile), with a flow rate of 150 nL/min. MS acquisition was performed in diaPASEF mode on the precursor mass range [400-1000] m/z, fragment mass range [100-1700] m/z, and ion mobility 1/K0 [0.64-1.37]. The acquisition scheme was composed of 8 consecutive TIMS ramps using an accumulation time of 100 ms, with 3 MS/MS acquisition windows of 25 Th for each of them. The resulting cycle time was 0.96 s. The collision energy was ramped linearly as a function of the ion mobility from 59 eV at 1/K0=1.6Vs cm−2 to 20 eV at 1/K0=0.6Vs cm−2. These files are available alongside all the associated metadata39 on the ProteomeXchange^40^ repository PRIDE^41^ with the following identifier: PXD062685.

#### Astral

Samples were analyzed in triplicate (50 ng) by nanoLC-MS/MS using a Vanquish Neo LC instrument (Thermo Fisher Scientific) coupled to a Thermo Orbitrap Astral (Thermo Fisher Scientific). Peptides were separated on a 50 cm μPAC™ column (Thermo Fisher Scientific), featuring a structured pillar array bed with a 180 µm bed width. Chromatographic separation was performed using a binary gradient system consisting of solvent A (0.1% formic acid in water) and solvent B (0.1% formic acid in 80% acetonitrile). The gradient profile began with initial conditions of 96% A and 4% B maintained for 1 min at 750 nL/min. The flow rate was then reduced to 250 nL/min, and a linear gradient was applied from 4% to 40% B over 15 min, followed by a rapid increase to 100% B over 1 min. The column was washed with 100% B for 2 min before returning to initial conditions (4% B) with a 7 min equilibration period. Full MS scans were acquired over a mass range of 380–980 m/z with detection in the Orbitrap at a resolution of 240,000. For DIA fragmentation, each acquisition cycle employed 300 isolation windows of 2 Th width, systematically covering the entire precursor ion range from 380 to 980 m/z42. All precursor ions within each window were fragmented using higher-energy collisional dissociation (HCD) with a normalized collision energy of 25%. MS2 spectra were acquired over a mass range of 150–2000 m/z with detection in the Astral mass analyzer, using a maximum injection time of 3 ms to optimize sensitivity while maintaining acquisition speed. The Astral .raw files are available alongside all the associated metadata^39^ on the ProteomeXchange^40^ repository PRIDE^41^ with the following identifier: PXD070049.

#### FASTA

The FASTA file for the DDA and DIA module was created by combining all UniProtKB/Swiss-Prot^19^ proteins for *Homo sapiens* (20,537 proteins), *Escherichia coli* (strain K12) (4,401 proteins), and *Saccharomyces cerevisiae* (6,722 proteins). The universal contaminant database^20^ and Biognosys iRT peptide sequences were also added.

## DATA AVAILABILITY

ProteoBench code, and all discussions are publicly available on the github repository github.com/Proteobench/ProteoBench. The full documentation (for users and contributors) is available here: proteobench.readthedocs.io. The files generated with the timsTOF SCP (Bruker) for the module of DIA quantification at precursor ion level on diaPASEF and Astral data are available alongside all the associated metadata^39^ on the ProteomeXchange^40^ repository PRIDE^41^ with the following identifier: PXD062685 and PXD070049, respectively. The data and Jupyter notebooks that were used to generate the figures of this manuscript are all publicly available on Zenodo with the following DOI: 10.5281/zenodo.17830658.

## CONTRIBUTION OF THE AUTHORS

Robbe Devreese (Software, Formal analysis, Investigation, Writing - Original Draft, Visualization), Caroline Jachmann (Software, Formal analysis, Investigation, Writing - Original Draft, Visualization), Bart Van Puyvelde (Investigation, Data Curation, Validation, Resources, Writing - Review & Editing), Holda A. Anagho-Mattanovich (Investigation, Writing - Original Draft, Funding acquisition), Witold E. Wolski (Software, Writing - Original Draft, Funding acquisition), Henry Webel (Software, Writing- Review & Editing), Matthias Anagho-Mattanovich (Software, Visualization, Funding acquisition), Wout Bittremieux (Methodology, Writing - Review & Editing, Funding acquisition), Karima Chaoui (Investigation, Data Curation, Validation, Writing - Review & Editing), Cristina Chiva (Investigation, Methodology, Writing - Review & Editing), Tine Claeys (Writing - Review & Editing), Hector Mauricio Castaneda Cortes (Software), Simon Devos (Funding Acquisition, Writing - Review & Editing), Maarten Dhaenens (Validation, Resources, Data Curation), Nadezhda T. Doncheva (Software, Writing Review & Editing), Viktoria Dorfer (Conceptualization, Methodology, Software, Writing - Review & Editing), Martin Eisenacher (Funding acquisition, Resources, Supervision, Writing - Review & Editing), Ralf Gabriels (Conceptualization, Methodology, Software, Writing - Review & Editing), Quentin Giai Gianetto (Formal analysis, Investigation, Writing - Review & Editing), David M. Hollenstein (Writing - Review & Editing, Software), Lars Juhl Jensen (Supervision, Funding Acquisition, Writing - Review & Editing), Vedran Kasalica (Software), Olivier Langella (Software, Writing - Review & Editing), Caroline Lennartsson (Software, Writing - Review & Editing), Dominik Lux (Writing - Review & Editing, Software), Lennart Martens (Supervision, Funding Acquisition, Writing - Review & Editing), Mariette Matondo (Writing - Review & editing), Teresa Mendes Maia (Writing - Review & editing), Emmanuelle Mouton-Barbosa (Data Curation, Validation, Writing - Review & Editing), Alireza Nameni (Software, Formal analysis, Investigation, Writing - Review & Editing), Michael Lund Nielsen (Supervision, Funding Acquisition, Writing - Review & Editing), Jesper Velgaard Olsen (Supervision, Funding acquisition, Writing - Review & editing), Magnus Palmblad (Validation, Writing - Review & editing, Funding acquisition), Christian Panse (Writing - Review & editing, Funding acquisition), Yasset Perez-Riverol (Writing - Review & editing, Software), Marina Pominova (Software), Martin Rykær (Conceptualization, Formal Analysis, Methodology, Writing – Review & editing), Eduard Sabidó (Data Acquisition, Methodology, Writing - Review & Editing), Julia Schessner (Software), Martin Schneider (Writing - Review & Editing), Veit Schwämmle (Software, Conceptualization, Formal analysis), An Staes (Writing - Review & editing), Maximilian T. Strauss (Conceptualization), Tim Van Den Bossche (Conceptualization, Writing - Review & Editing), Sam van Puyenbroeck (Software, Methodology), Kevin Velghe (Software), Runxuan Zhang (Writing - Review & Editing), Julian Uszkoreit (Software, Writing - Review & Editing, Resources, Project administration), Robbin Bouwmeester (Supervision, Conceptualization, Software, Writing - Original Draft, Visualization, Project administration), Marie Locard-Paulet (Supervision, Conceptualization, Software, Writing - Original Draft, Visualization, Project administration).

## ACKNOWLEDGEMENTS

We would like to acknowledge the European Bioinformatics Community for Mass Spectrometry (EuBIC-MS), an initiative of the European Proteomics Association (EuPA), for supporting this project. ProteoBench is hosted and supported as an infrastructure service of the Core Unit Bioinformatics at the Medical Faculty of the Ruhr University Bochum - CUBiMed.RUB. This work was also supported by the de.NBI Cloud within the German Network for Bioinformatics Infrastructure (de.NBI) and ELIXIR-DE (Forschungszentrum Jülich and W-de.NBI-001, W-de.NBI-004, W-de.NBI-005 W-de.NBI-008, W-de.NBI-010, W-de.NBI-013, W-de.NBI-014, W-de.NBI-016, W-de.NBI-022). The Danish Data Science Academy (DDSA-SE-2022-017), FWO (W000924N), and the Core for Life consortium (coreforlife.sites.vib.be) funded our in-person meetings. We also want to thank all the principal investigators who gave their students, postdocs, and collaborators the funding and freedom to contribute to ProteoBench. Finally, we want to thank all the people who spent time discussing how to best tackle the challenge of comparing proteomics data analysis workflows.

M.L.-P., E.M.-B. and K.C. were supported by grants from the French National Research Agency (ProFI projects: ANR-10-INBS-08 & ANR-24-INBS-0015), the Région Occitanie, and the REACT-EU program of the European Commission. R.D., B.V.P., R.G., L.M., R.B., W.B., and T.V.D.B. acknowledge funding from the Research Foundation Flanders (FWO) [1SH9O24N, 1278023N, 12AK526N, G010023N, G028821N, 12A6L24N, G087625N, 1286824N]. A.N. acknowledges funding from the European Union’s Horizon 2020 research and innovation programme under the Marie Skłodowska-Curie grant agreement no 956148. C.J. acknowledges funding from the European Union’s Horizon Europe under grant agreement no. 101119980. L.M. acknowledges funding from the Horizon Europe Projects BAXERNA 2.0 [101080544] and COMBINE [101191739], and from the Ghent University Concerted Research Action [BOF21/GOA/033]. L.M. is further supported by the CHIST-ERA project ODEEP-EU [G0GDV23N]. V.K. and M.Pa. acknowledge funding from the NLeSC Open eScience grant number 27021G10. M.A.-M. was supported by the Novo Nordisk foundation with an unconditional donation to the NNF Center for Basic Metabolic research with the grant number NNF23SA0084103. Work at The Novo Nordisk Foundation Center for Protein Research (CPR) is funded in part by a donation from the Novo Nordisk Foundation (NNF14CC0001, NNF24SA0098829, NNF13OC0006477 and NNF21OC0072070). This project was supported by a center-of-excellence grant from the Danish National Research Foundation to Copenhagen Center for Glycocalyx Research (DNRF196). This project was also supported by a grant from the Danish Agency of Higher Education and Science to establish the PLATO research infrastructure: Danish National Mass Spectrometry Platform for Proteomics and Biomolecular Imaging (5229-00012B). J.V.O. is also funded by Novo Nordisk a/s (CELFFI-2022-002843). H.A.A.-M. additionally acknowledges funding from the NNF Bioscience PhD Grant NNF19SA0035440. W.B. and M.Po. acknowledge support from the University of Antwerp’s University Research Fund. H.M.C.C and R.Z. acknowledge funding from the Scottish Government RESAS project KJHI-B1-2 and BBSRC project BB/Y513192/1. D.L. acknowledges funding from the InnovationsFoRUM research funding of the Medical Faculty, Ruhr-University Bochum (ProMiS, IF-014N-22). M.E. was partly funded by the German Federal Ministry of Research, Technology and Space (BMFTR) in the frame of de.NBI/ELIXIR-DE (W-de.NBI-005). E.S. acknowledges support of the Spanish Ministry of Science and Innovation through the Centro de Excelencia Severo Ochoa (CEX2020-001049-S grant funded by MCIN/AEI/10.13039/501100011033) and PID2020-115092GB-I00 funded by AEI/10.13039/501100011033, and the Generalitat de Catalunya through the CERCA programme and the Departament de Recerca i Universitats (2021-SGR2021-01225). The CRG/UPF Proteomics Unit is part of the Spanish Infrastructure for Omics Technologies (ICTS OmicsTech).

