## Supplementary Figure 1 for "ProteoBench: the community-curated platform for comparing proteomics data analysis workflows"

Home

› DDA

› DIA

› Archived

privacy notice

legal notice

👈 Select a page from the sidebar to get started!
📖 Learn more about Proteobench on proteobench.readthedocs.io
💻 Find the source code on github.com

ProteoBench is a project initiated and maintained by EuBIC-MS. Please join us and contribute!

If you still have questions, you can email us here

Using proteobench version: 0.10.4

This site is hosted by the Core Unit Bioinformatics at the Medical Faculty of the Ruhr University Bochum - CUBiMed.RUB (https://cubimed.rub.de), which is co-funded by the BMBF-funded German Network for Bioinformatics Infrastructure (de.NBI)

### ProteoBench Overview

Active modules

**8**

Proposed and in-development modules

**6**

Supported workflows and tools

**16**

Submitted points

**204**

Monthly visitors

**213**
