## Supplementary Figure 2 for "ProteoBench: the community-curated platform for comparing proteomics data analysis workflows"

### DIA Precursor ion quantification - Astral 2 Th

Go to module documentation

This module is in BETA phase. The figure presented below and the metrics calculation may change in the near future.

### Results (All Data)

Choose with the slider below the minimum number of quantification value per raw file.
Example: when 3 is selected, only the precursor ions quantified in 3 or more raw files will be considered for the plot.

Minimal precursor quantifications (# samples)

3

1 6

Select label to plot

None

Select metric to plot

Median
Mean

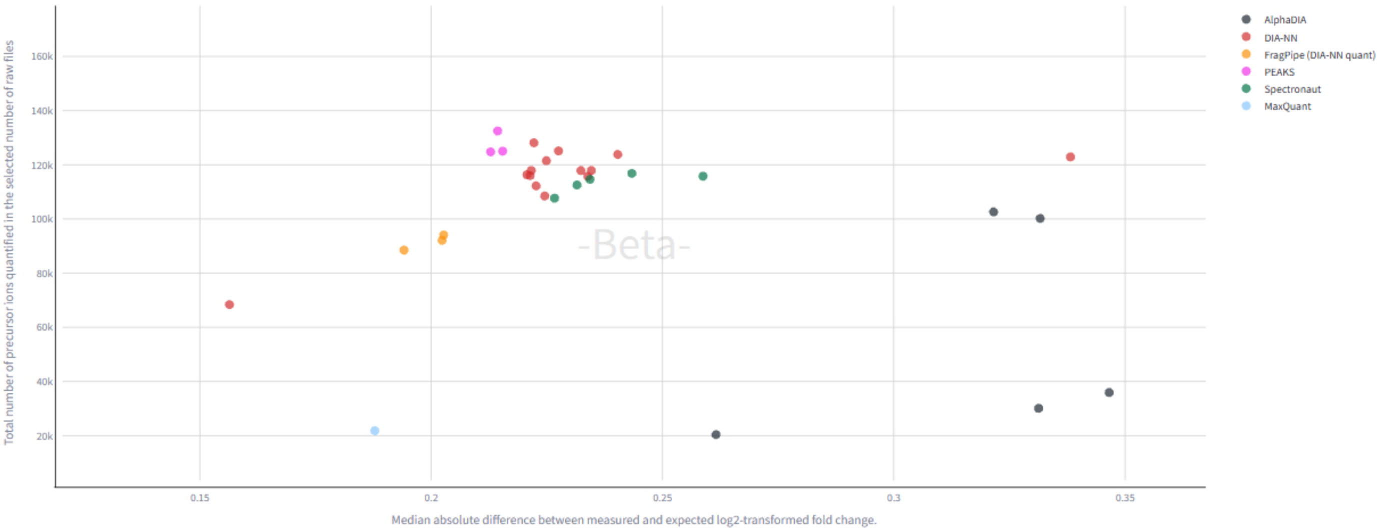


| | old_new | id | software_name | software_version | search_engine | search_engine_version | ident_fdr_psm | ident_fdr_peptide | ident_fdr_protein | enable_match_between_runs | precursor_mass_t |
|---|---|---|---|---|---|---|---|---|---|---|---|
| 0 | old | AlphaDIA_20250605_164604 | AlphaDIA | 1.10.3 | AlphaDIA | None | 0.01 | None | 0.01 | True | [-3 ppm, 3 ppm] |
| 1 | old | DIA-NN_20250605_175300 | DIA-NN | 2.0 Academia | DIA-NN | 2.0 Academia | 0.01 | None | None | True | [-2.6 ppm, 2.6 ppr |
| 2 | old | FragPipe (DIA-NN quant)_20250704_132452 | FragPipe (DIA-NN | 23.0 | MSFragger | 4.3 | 0.01 | None | 0.01 | False | [-20 ppm, 20 ppm |
| 3 | old | DIA-NN_20250605_165130 | DIA-NN | 2.1.0 Academia | DIA-NN | 2.1.0 Academia | 0.01 | None | None | True | [-10.0 ppm, 10.0 p |
| 4 | old | DIA-NN_20250605_165839 | DIA-NN | 1.7.16 | DIA-NN | 1.7.16 | 0.01 | None | None | True | [-2.78001 ppm, 2.7 |
| 5 | old | DIA-NN_20250626_092915 | DIA-NN | 1.9.2 | DIA-NN | 1.9.2 | 0.01 | None | 0.01 | True | [-2.31639 ppm, 2.3 |
| 6 | old | DIA-NN_20250605_170158 | DIA-NN | 1.9.2 | DIA-NN | 1.9.2 | 0.01 | None | None | True | [-2.42372 ppm, 2.4 |
| 7 | old | DIA-NN_20250704_085203 | DIA-NN | 2.2.0 Academia | DIA-NN | 2.2.0 Academia | 0.01 | None | None | True | [-2.6 ppm, 2.6 ppr |
| 8 | old | AlphaDIA_20250627_150126 | AlphaDIA | 1.10.4-dev0 | AlphaDIA | None | 0.01 | None | 0.01 | True | 0 |
| 9 | old | AlphaDIA_20250605_170349 | AlphaDIA | 1.10.3 | AlphaDIA | None | 0.01 | None | 0.01 | False | [-10 ppm, 10 ppm |

#### Download raw datasets

Select dataset

Choose an option
