## Supplementary Figure 3 for "ProteoBench: the community-curated platform for comparing proteomics data analysis workflows"

Public Benchmark Runs | Submit New Data | Results In-Depth | Results New Data | Public Submission

### DIA Precursor ion quantification - Astral 2 Th

Go to module documentation

This module is in BETA phase. The figure presented below and the metrics calculation may change in the near future.

#### Input files

Please upload the output of your analysis, and indicate what software tool it comes from (this is necessary to correctly parse your table) - find more information in the "How to use" section of this module. To the raw data to run your workflow on please see the section "Data set".

Remember: contaminant sequences are already present in the fasta file associated to this module. **Do not add other contaminants** to your search. This is important when using MaxQuant and FragPipe, among other tools.

Software tool

DIA-NN

Software tool result file

Drag and drop file here
Limit 1GB per file

Browse files

report.parquet 68.7MB

Now, press `Parse and Bench` to calculate the metrics from your input.

Parse and bench
