## Supplementary Figure 4 for "ProteoBench: the community-curated platform for comparing proteomics data analysis workflows"

### DDA Precursor quantification (QExactive)

Go to module documentation

This module is in BETA phase. The figure presented below and the metrics calculation may change in the near future.

#### Select dataset to plot

Select dataset

MaxQuant_20250605_122203

#### Log2 Fold Change distributions by species.

log2 fold changes calculated from MaxQuant_20250605_122203

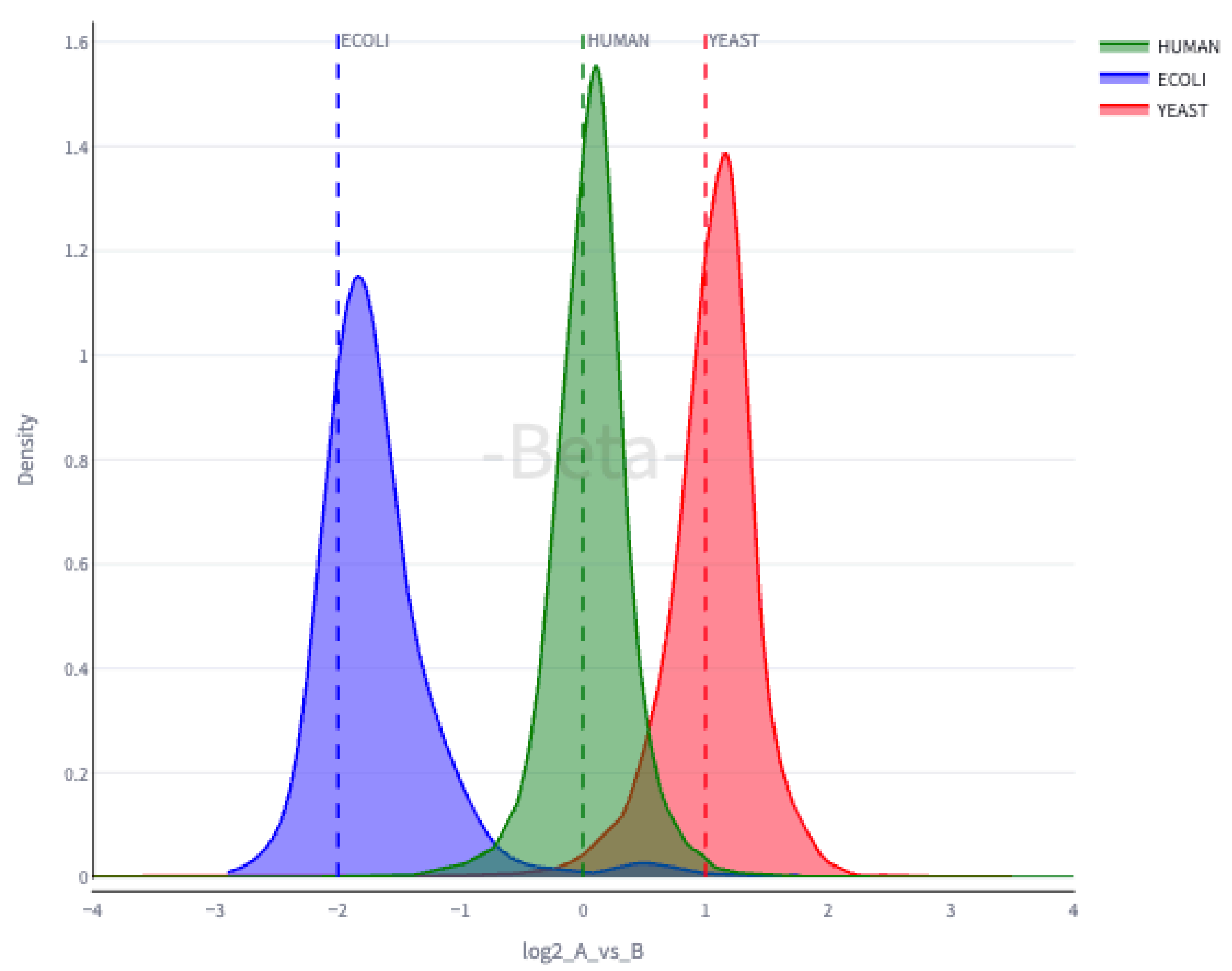


#### Coefficient of variation distribution in Condition A and B.

CVs calculated from MaxQuant_20250605_122203

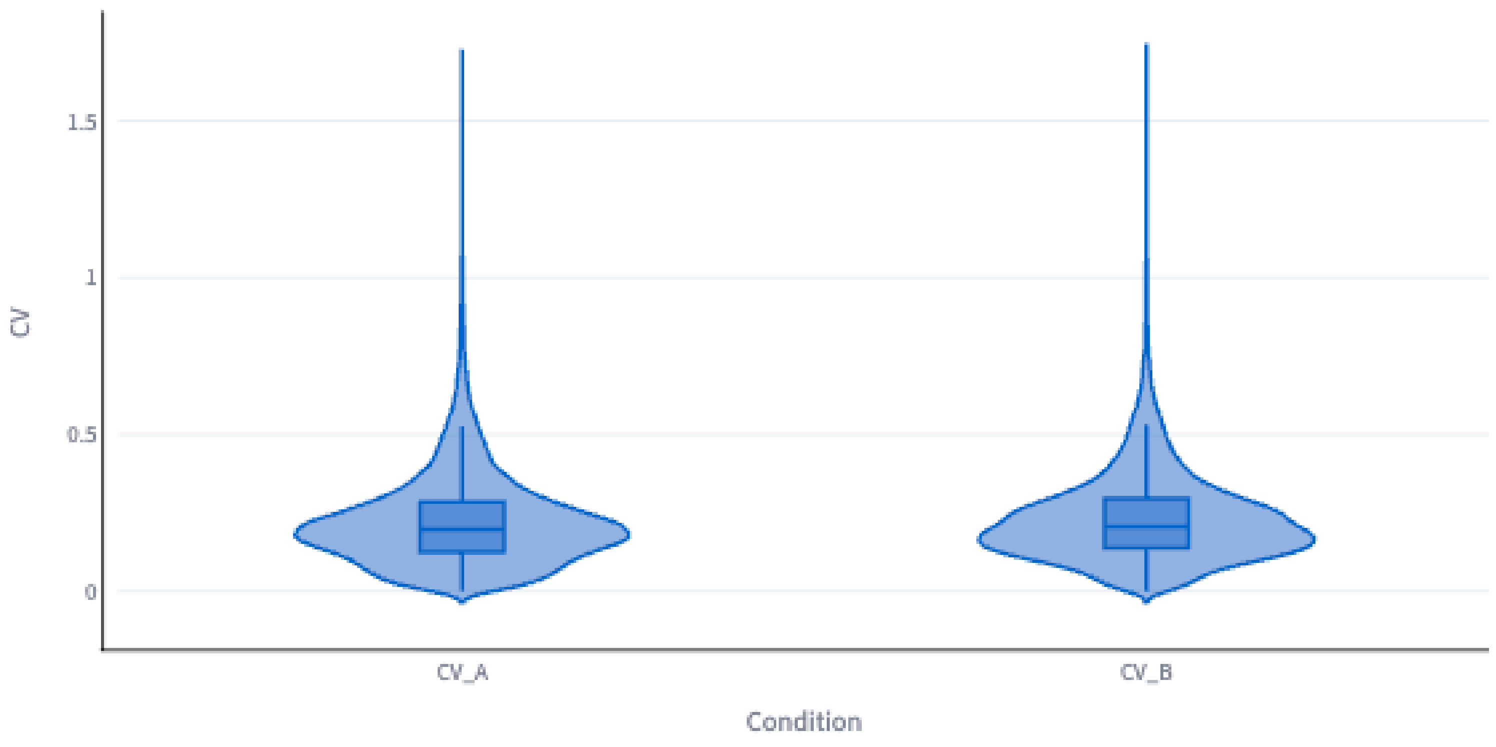


#### MA plot

MA plot calculated from MaxQuant_20250605_122203

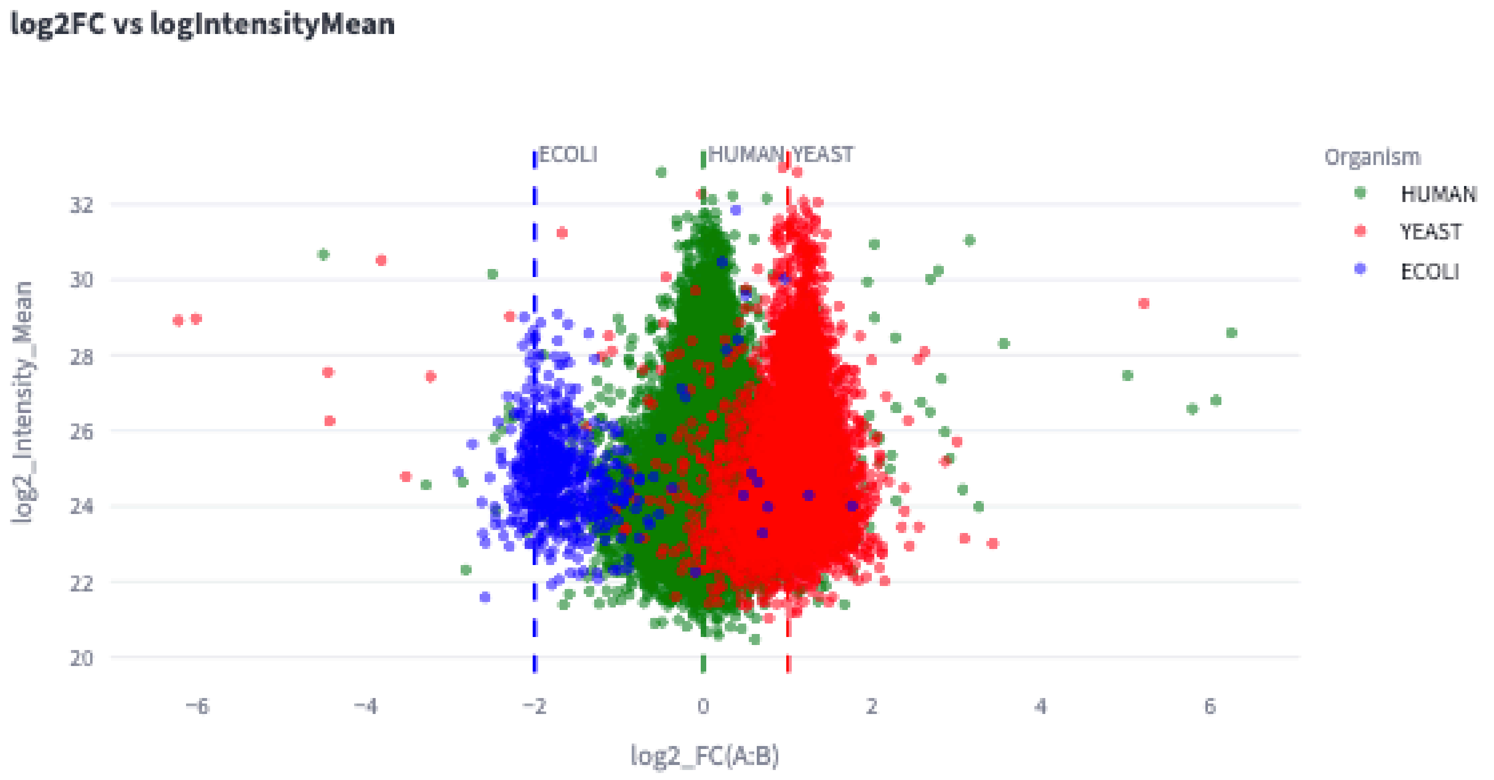


#### Sample of the processed file for MaxQuant_20250605_122203

Here are the results from your benchmark run. The table contains the precursor ion MS signal calculated from your input data. You can download this table from `Download calculated ratios` below or via the download button in the top right corner.

| | precursor ion | log_Intensity_mean_A | log_Intensity_mean_B | log_Intensity_std_A | log_Intensity_std_B | Intensity_mean_A | Intensity_mean_B | Intensity_std_A | Intensity_std_B | CV_A | CV_B | log2_A |
|---|---|---|---|---|---|---|---|---|---|---|---|---|
| 5 | AAAASAAEAGIATTGTEDSDDALLK\|Z=3 | 25.2148 | 25.327 | 0.1053 | 0.1418 | 38993000 | 42227666.6667 | 2843983.4739 | 4143260.592 | 0.0729 | 0.0981 | -[illegible] |
| 7 | AAADALSDLEIK\|Z=2 | 27.0888 | 26.1368 | 0.4413 | 0.2297 | 146956666.6667 | 74392666.6667 | 40440537.006 | 11300504.7822 | 0.2752 | 0.1519 | [illegible] |
| 9 | AAAEDVNVTFEDQQK\|Z=2 | 27.0872 | 26.4451 | None | 0.4265 | 142580000 | 93935666.6667 | None | 25782597.8973 | None | 0.2745 | [illegible] |
| 10 | AAAEGVANLHLDEATGEMVSK\|Z=3 | 25.1165 | None | 0.166 | None | 36536333.3333 | None | 4116457.8625 | None | 0.1127 | None | |
| 13 | AAAEVAGQFVIK\|Z=2 | 26.9125 | 26.6334 | 0.0649 | 0.4641 | 126406666.6667 | 107611000 | 5724249.5869 | 32607424.9673 | 0.0453 | 0.303 | [illegible] |
| 14 | AAAEVNQDYGLDPK\|Z=2 | 27.4574 | 27.3389 | 0.283 | 0.4271 | 186576666.6667 | 174733333.3333 | 34423373.3578 | 50786299.6618 | 0.1845 | 0.2907 | [illegible] |
| 15 | AAAFEEQENETVVVK\|Z=2 | 25.4437 | 25.4335 | 0.3666 | 0.55 | 46587333.3333 | 47492333.3333 | 11060116.5154 | 17275383.2181 | 0.2374 | 0.3638 | [illegible] |
| 16 | AAAFEGELIPASQIDR\|Z=2 | 25.132 | 27.1381 | 0.3542 | 0.183 | 37481333.3333 | 148483333.3333 | 8572077.7722 | 18233788.2332 | 0.2287 | 0.1228 | -[illegible] |
| 19 | AAAGELQEDSGLC[Carbamidomethyl]VLAR\|Z= | 24.3634 | 24.6205 | 0.574 | None | 22443500 | 25793000 | 8700948.9425 | None | 0.3877 | None | -[illegible] |
| 20 | AAAGGQGSAVAAEAEPGKEEPPAR\|Z=3 | 23.8408 | 23.0595 | 0.4772 | 0.5556 | 15437000 | 9207466.6667 | 5014801.2922 | 3783941.3649 | 0.3249 | 0.411 | [illegible] |

#### Download table

Download
