## Supplementary Figure 6 for "ProteoBench: the community-curated platform for comparing proteomics data analysis workflows"

Public Benchmark Runs | Submit New Data | Results In-Depth | Results New Data | Public Submission

### DIA Precursor ion quantification - Astral 2 Th

Go to module documentation

This module is in BETA phase. The figure presented below and the metrics calculation may change in the near future.

Meta data for searches

Drag and drop files here
Limit 1GB per file

Browse files

report.log.txt 24.2KB

Additionally, you can fill out parameters for your search manually. Please, only fill out the parameters that are not already included in the input file. Only make changes if you are sure about the parameters you are changing.

Software name: DIA-NN

Software tool version: 2.2.0 Academia

Search engine name: DIA-NN

Search engine version: 2.2.0 Academia

FDR psm: 0.01

FDR peptide: None

FDR protein: None

- [x] Quantified with MBR

Precursor mass tolerance (including unit ppm, PPM or Da): [-2.5 ppm, 2.5 ppm]

Fragment mass tolerance (including unit ppm, PPM or Da): [-6 ppm, 6 ppm]

Proteolytic Enzyme: Trypsin/P

Maximum allowed number of missed cleavage: 1

Minimum peptide length: 6

Maximum peptide length: 30

Specify the fixed mods that were set: unimod4

Specify the variable mods that were set (separated by a comma): UniMod:35/15.994915/M,UniMod:1/42.010565/*n,UniMod:21/79.966331

Maximum number of modifications: 1

Minimum precursor charge allowed: 1

Maximum precursor charge allowed: 5

Quantification method: QuantUMS high-precision

Protein inference method: Genes

Abundance normalization method: None

Utilized spectral library: None

Window scanning size: None

Comments for submission

Anything else you want to let us know? Please specifically
add changes in your search parameters here, that are not obvious from the parameter file.

- [ ] I confirm that the metadata is correct
