## Supplementary figures and images for "ProteoBench: the community-curated platform for comparing proteomics data analysis workflows"

### Supplementary Figure 2

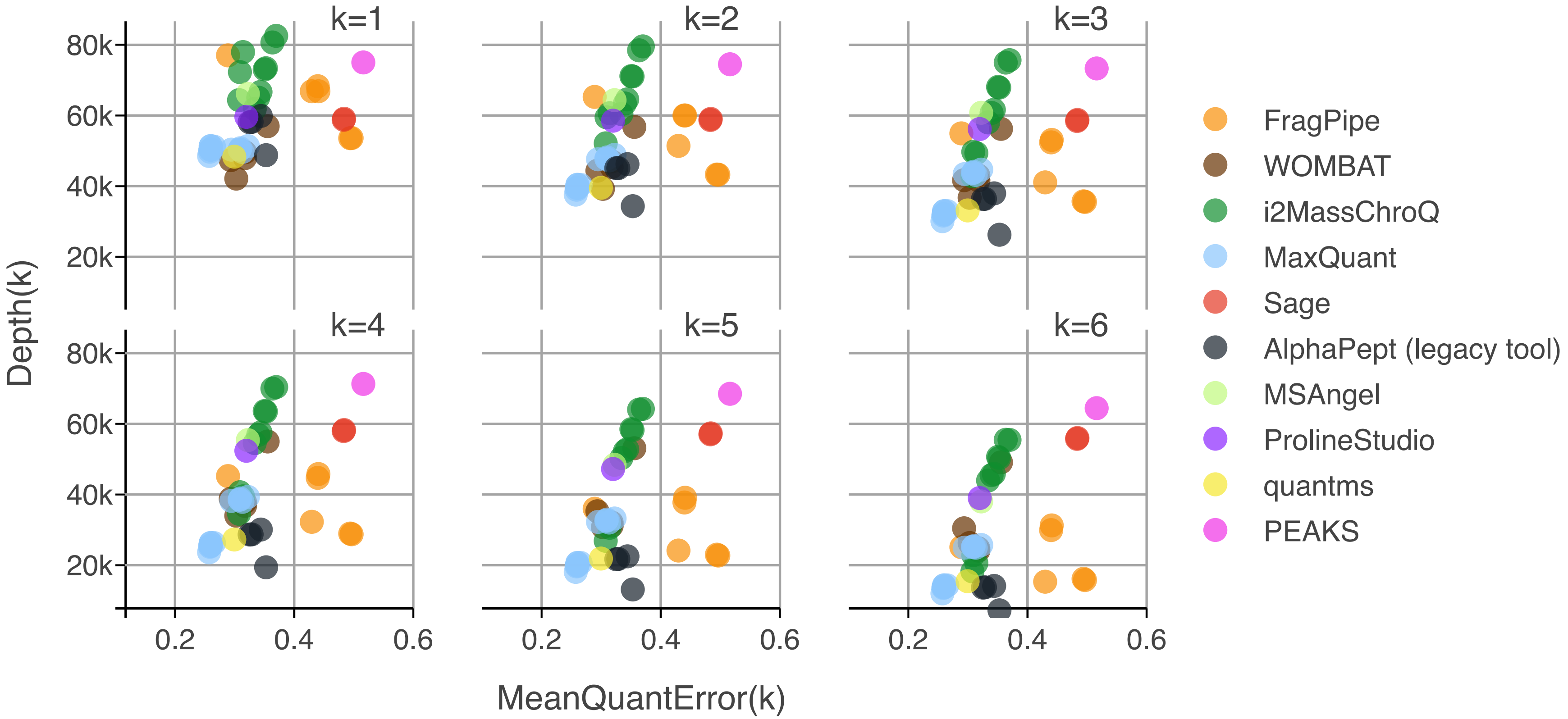

### Supplementary Figure 8

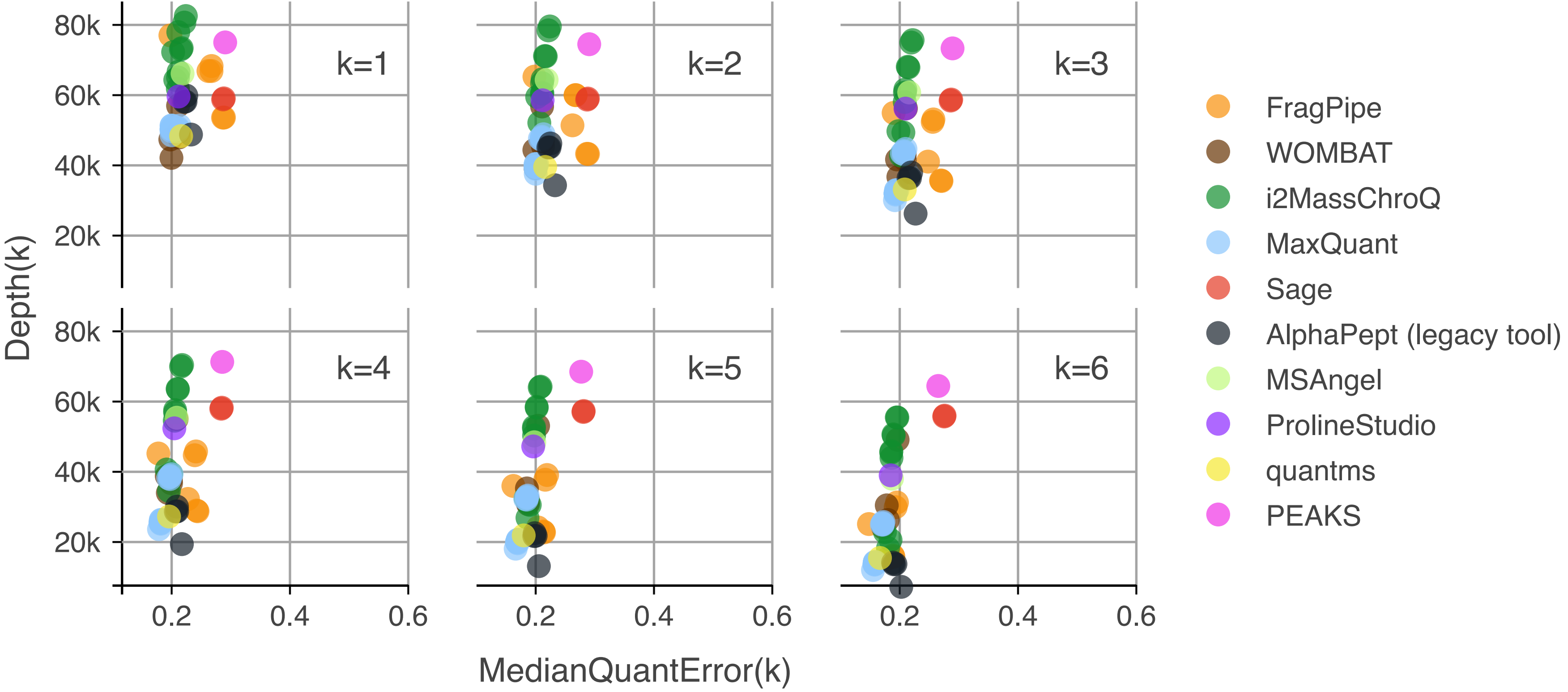

### Supplementary Figure 9

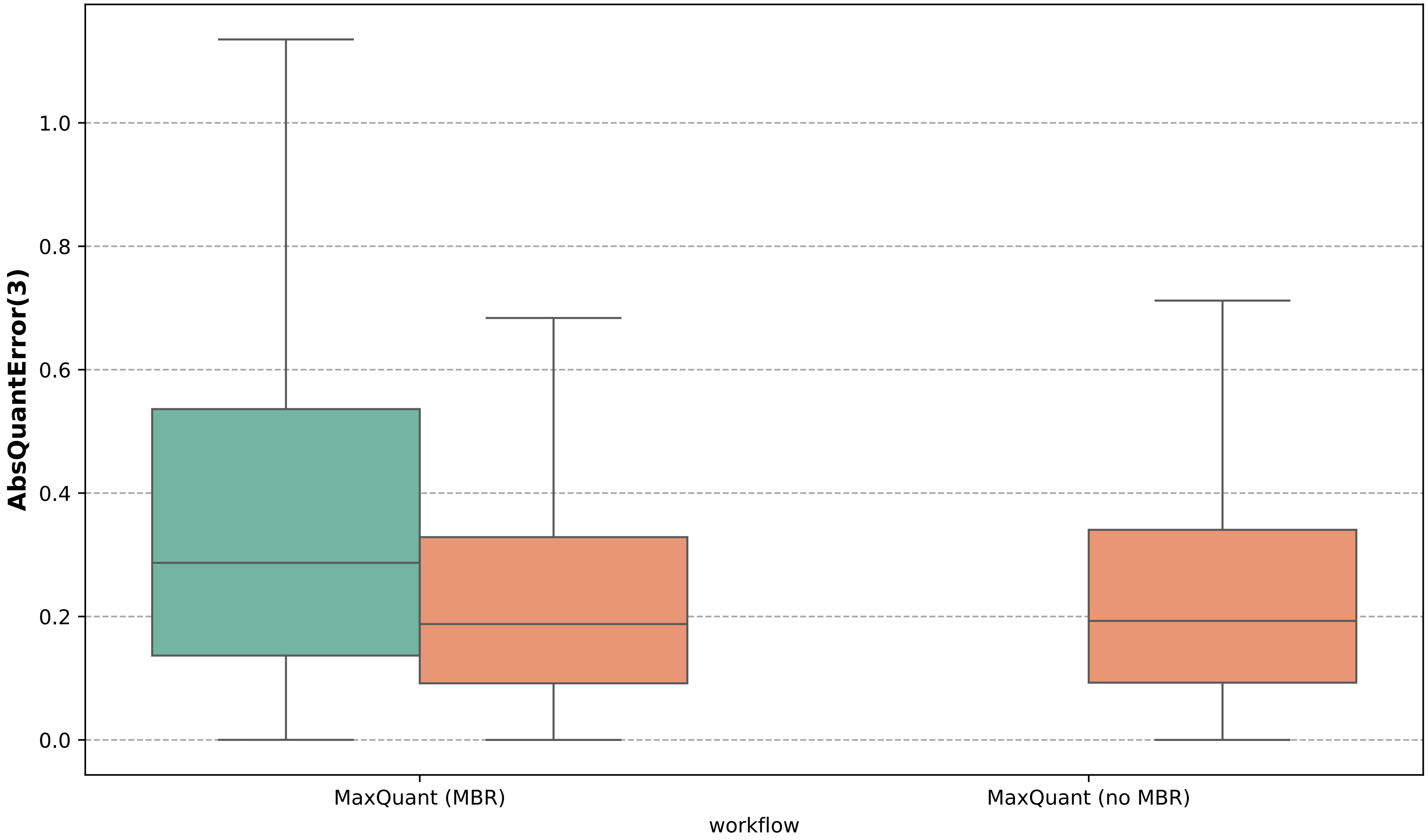

### Supplementary Figure 10

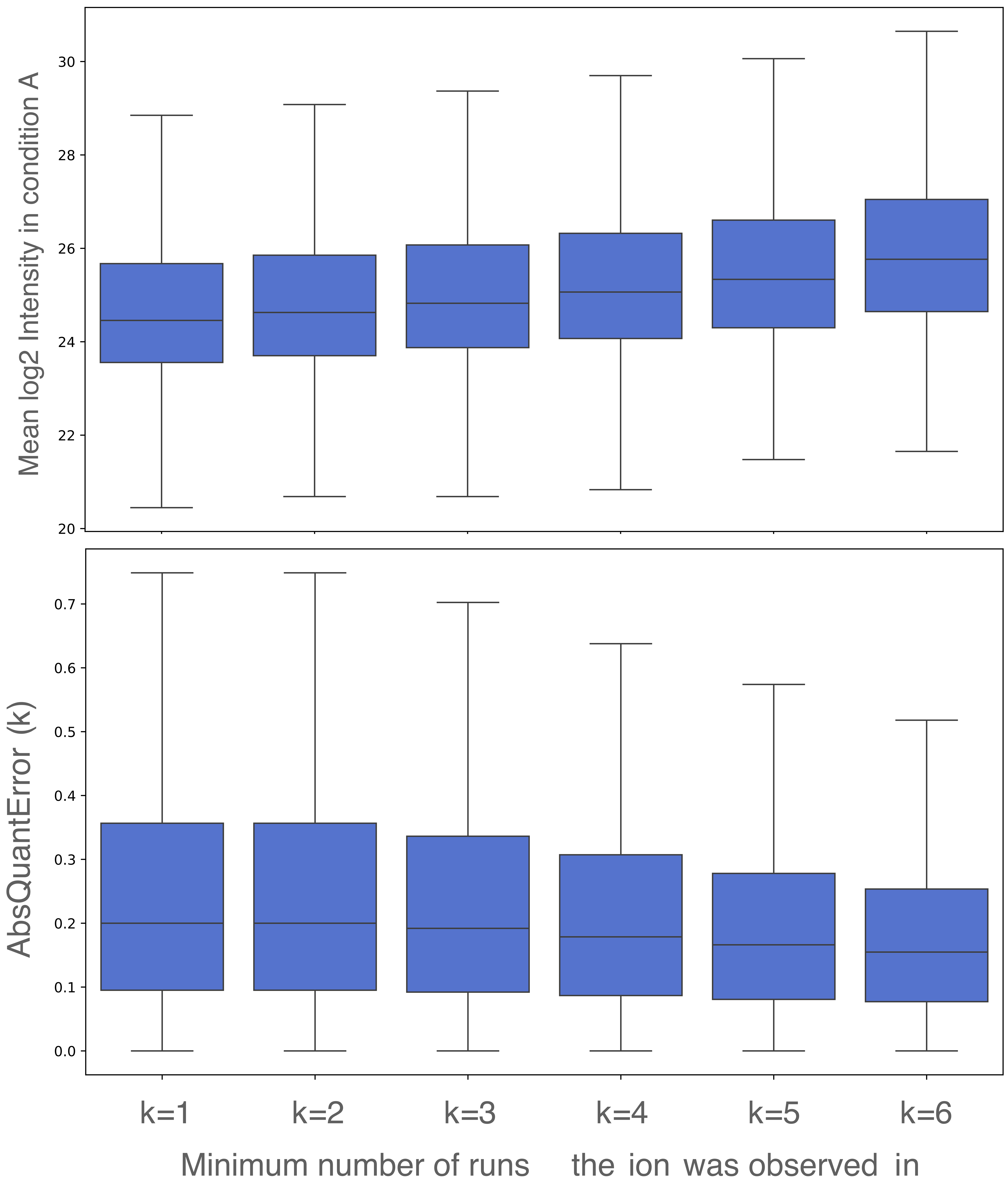

### Supplementary Figure 11

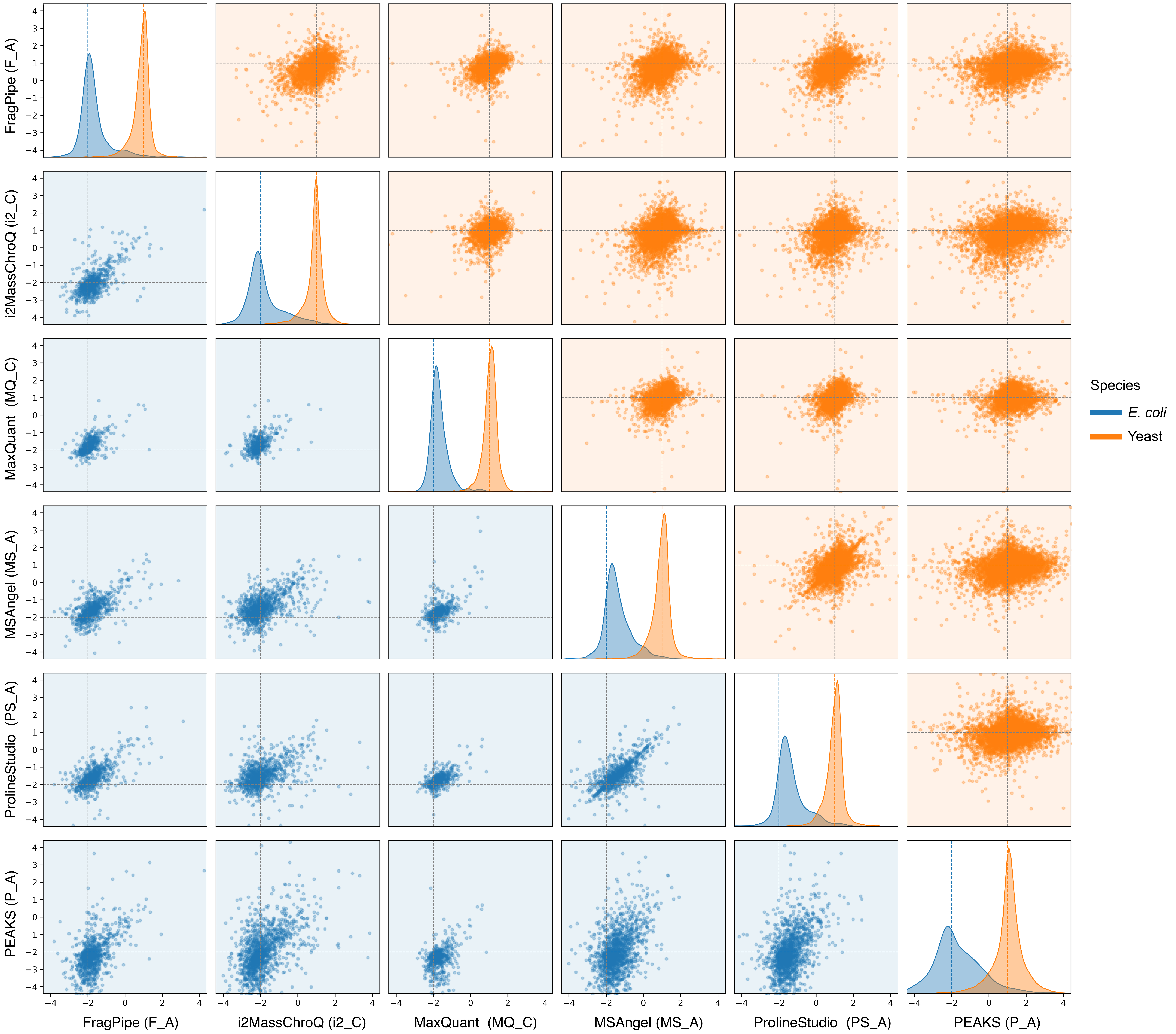

### Supplementary Figure 12

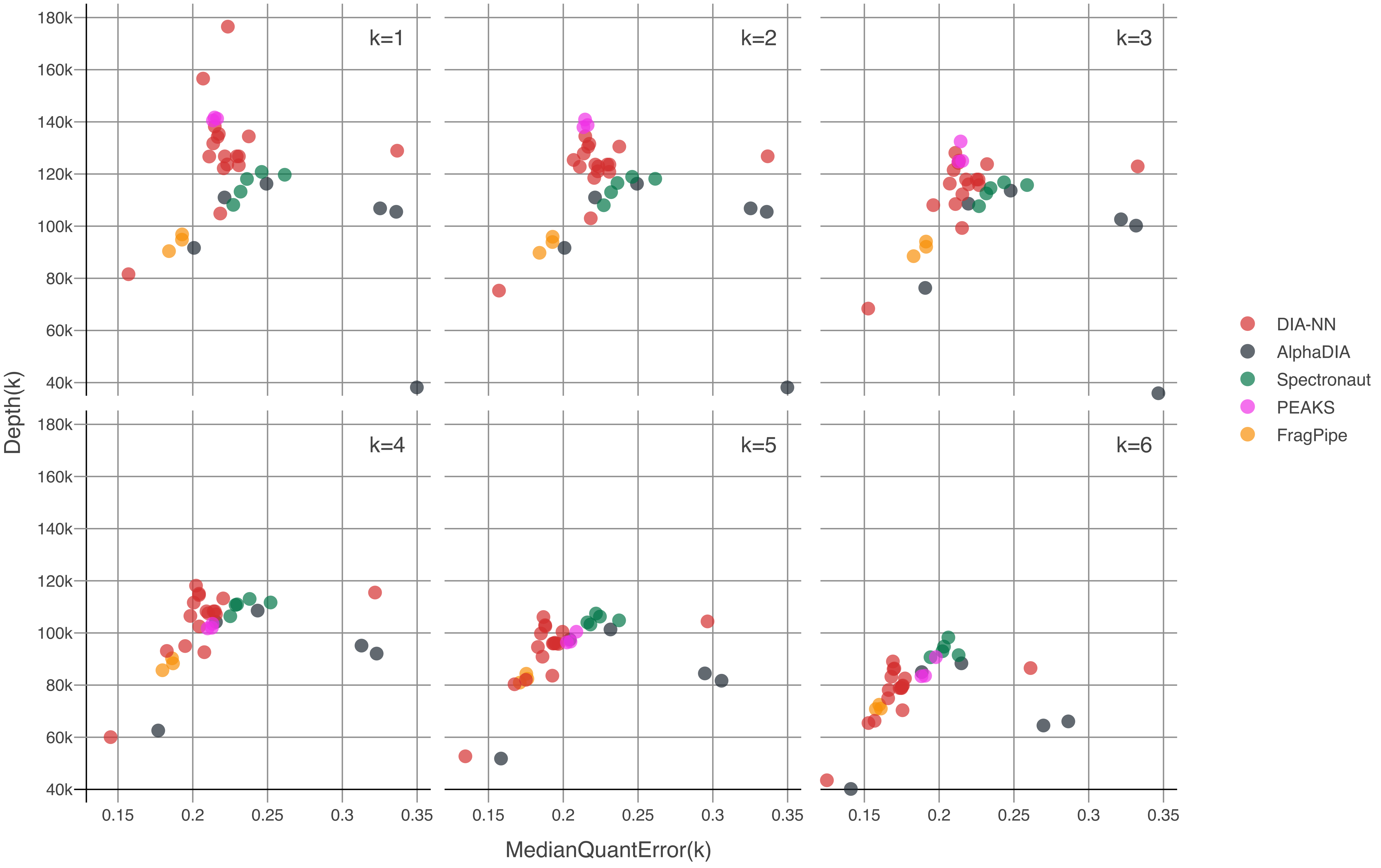

### Supplementary Figure 13

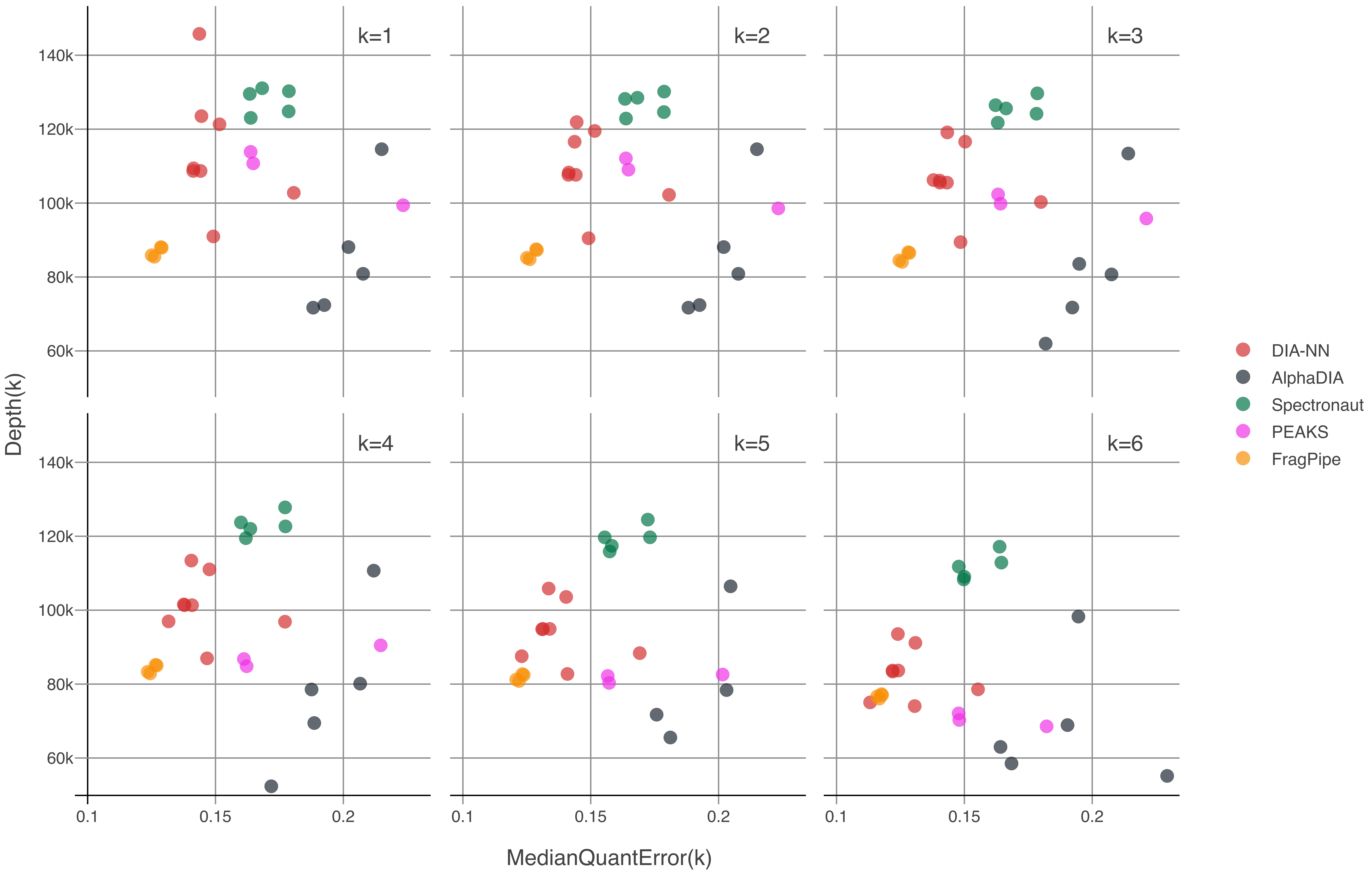

### Supplementary Figure 14

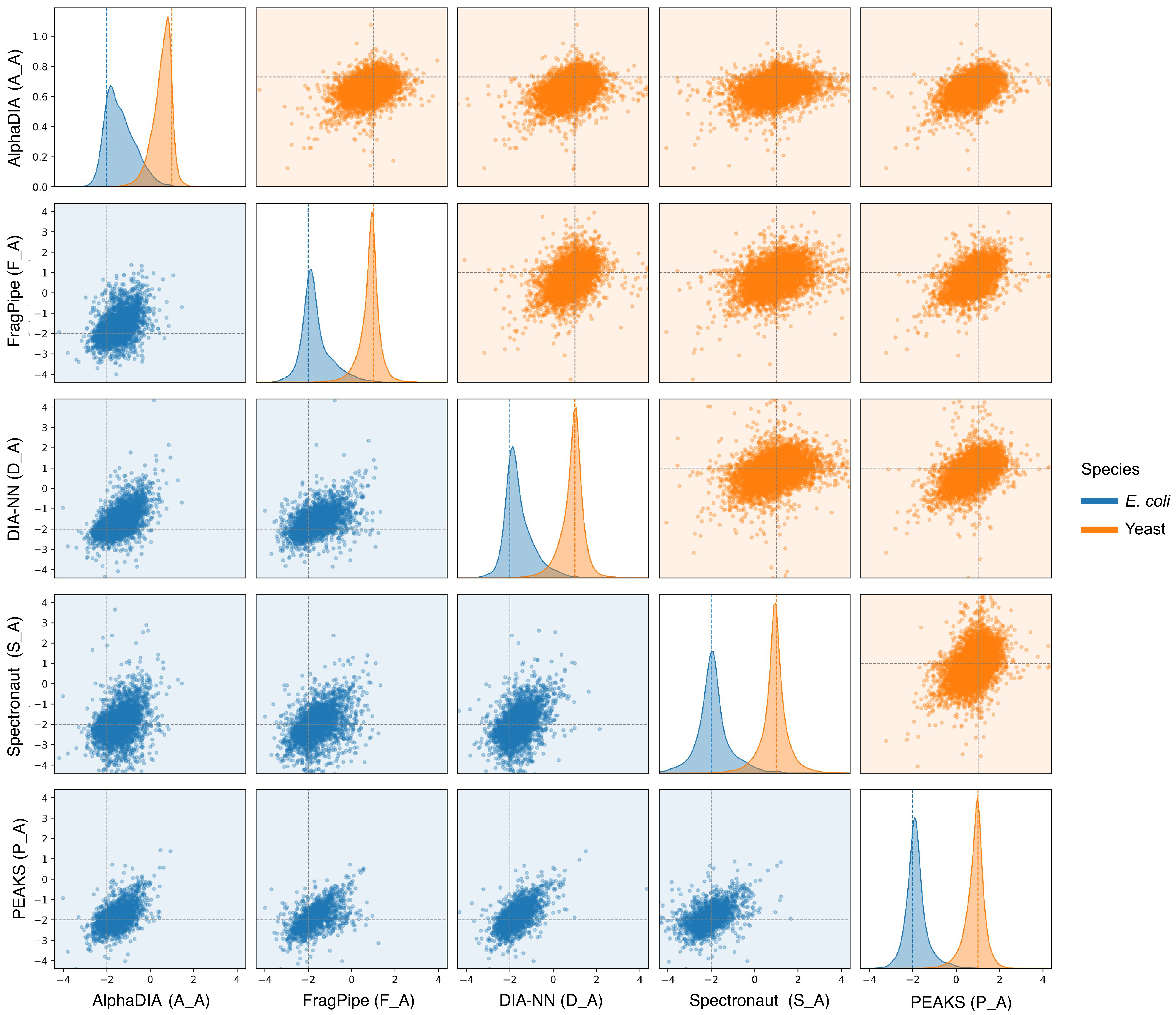
