## Supplementary Figures 1-14 for "ProteoBench: the community-curated platform for comparing proteomics data analysis workflows"

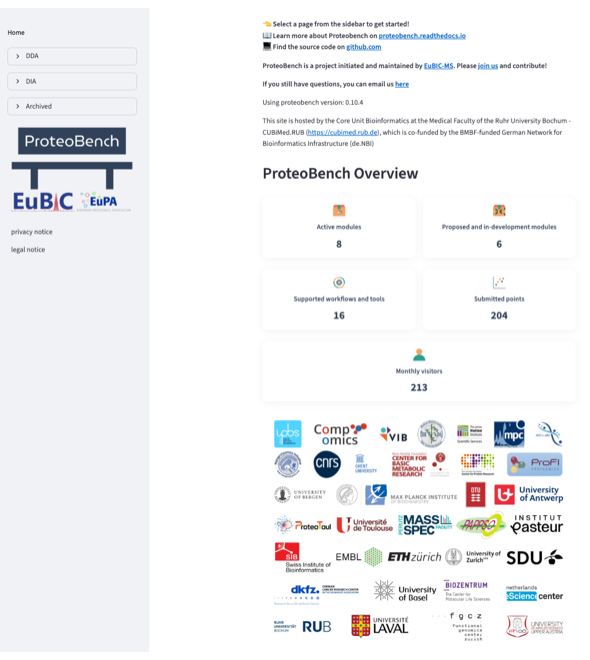

**Supplementary Figure 1:** **Home page of the ProteoBench user interface (proteobench.cubimed.rub.de)**.

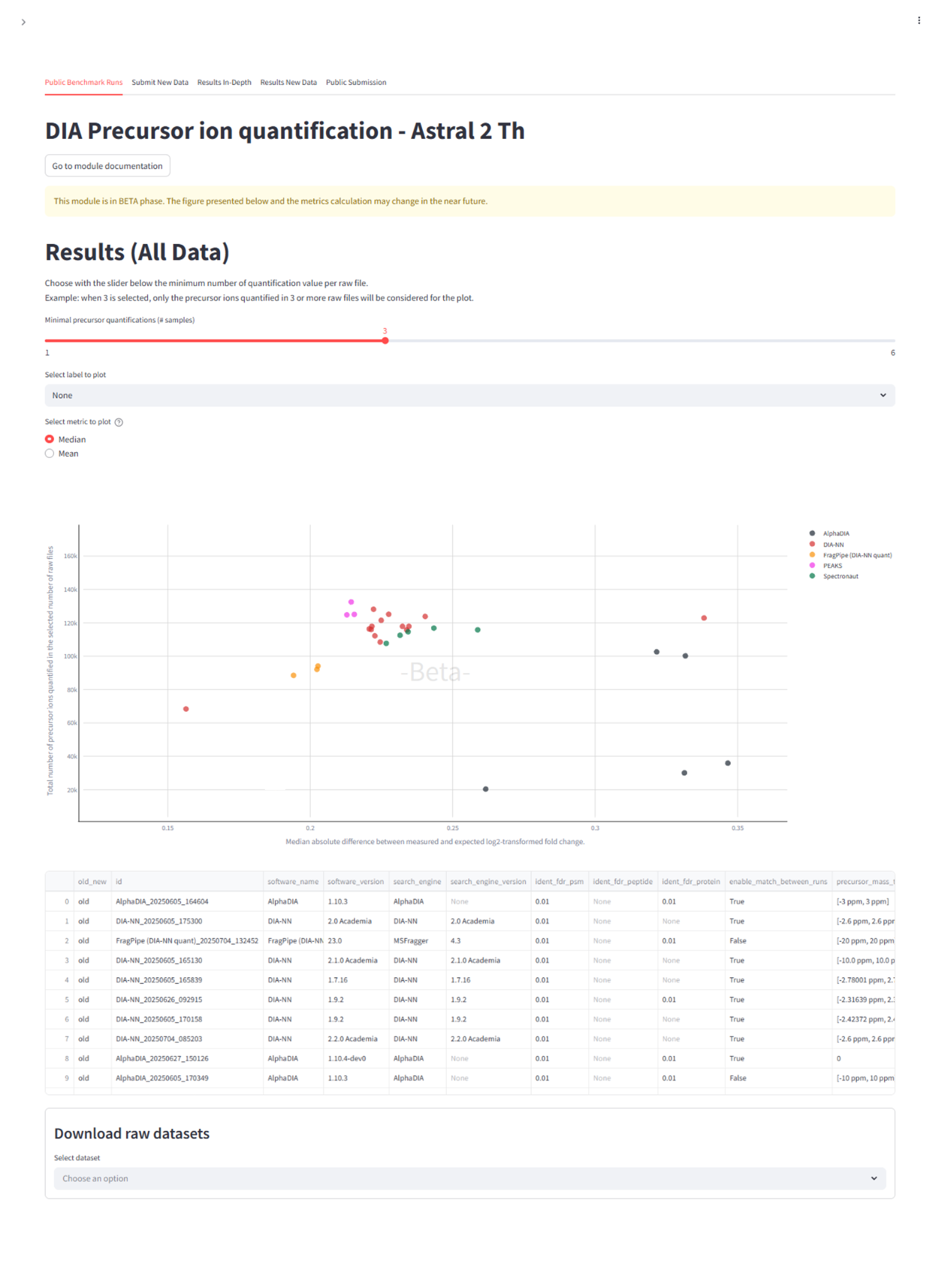

**Supplementary Figure 2:** **Main overview page of the Astral DIA precursor ion quantification module.**

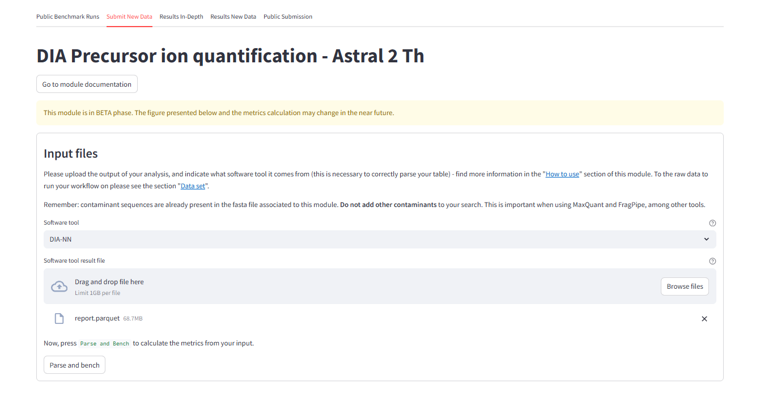
**Supplementary Figure 3: Upload tab of the Astral DIA precursor ion quantification module.**

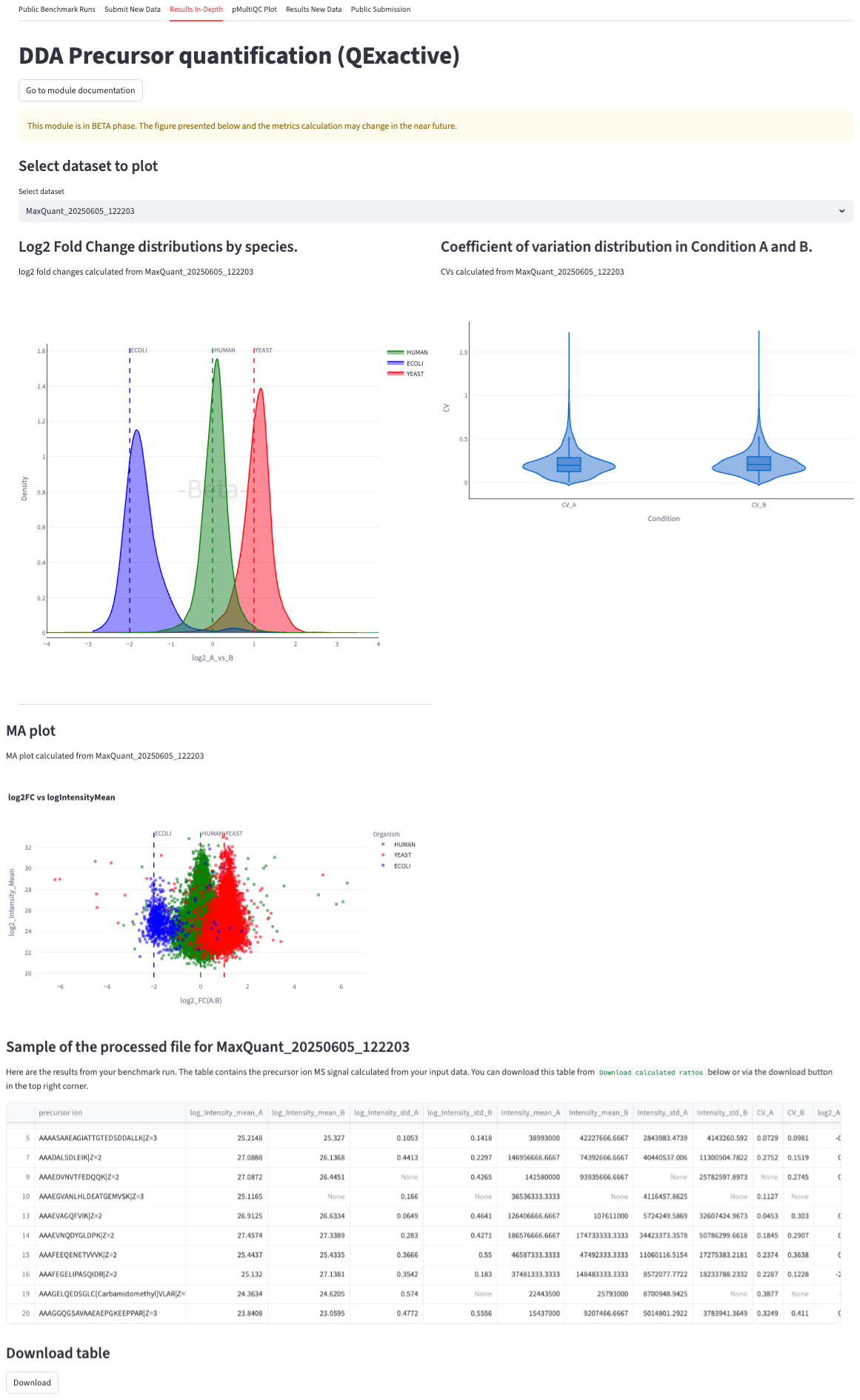

**Supplementary Figure 4:** **In-depth result tab for user-uploaded results of the module DDA precursor ion quantification module (Q Exactive).**

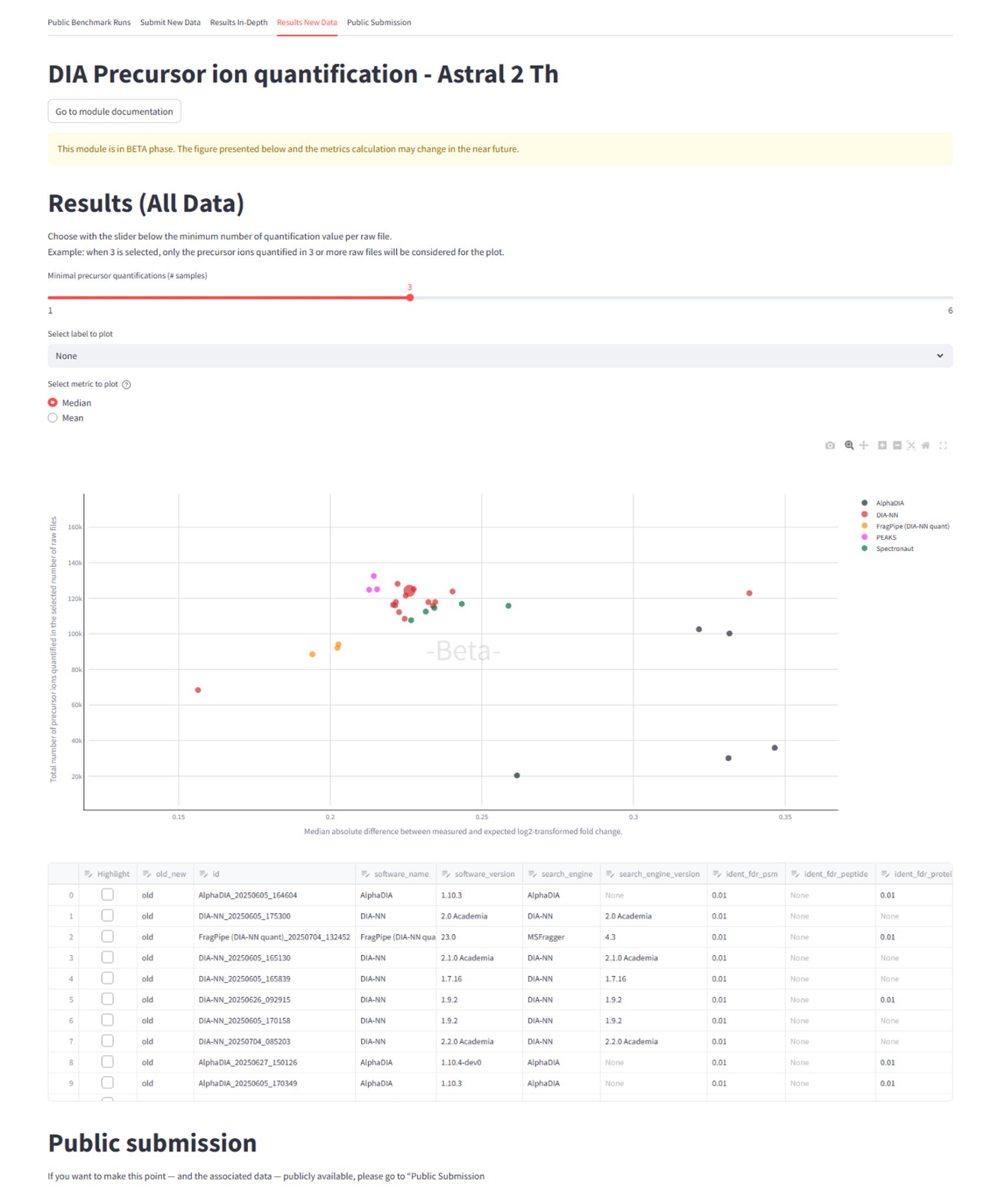

**Supplementary Figure 5: Result tab of the Astral DIA precursor ion quantification module.** It combines public points and user-uploaded points (larger point) in one figure.

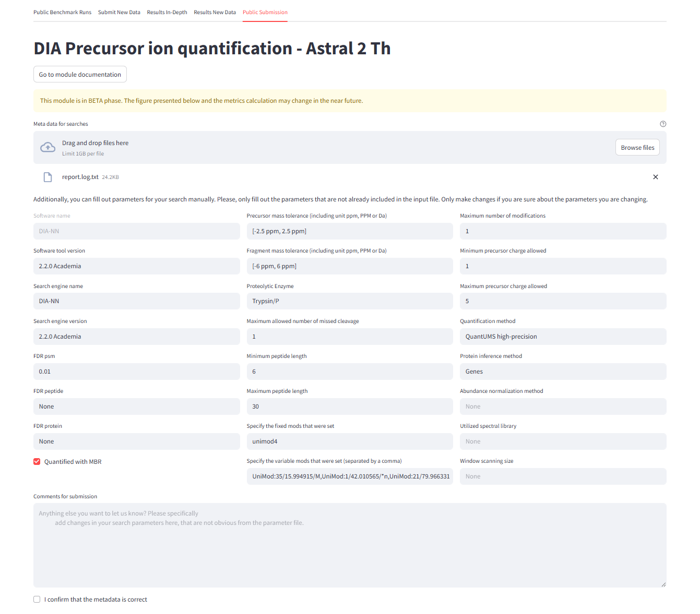

**Supplementary Figure 6: Public Submission tab of the Astral DIA precursor ion quantification module.** Parameters are parsed automatically from provided parameters or log files, and can be manually corrected if necessary. A free-text comment field is provided to allow the user to contribute further information about the workflow beyond the parsed parameters.

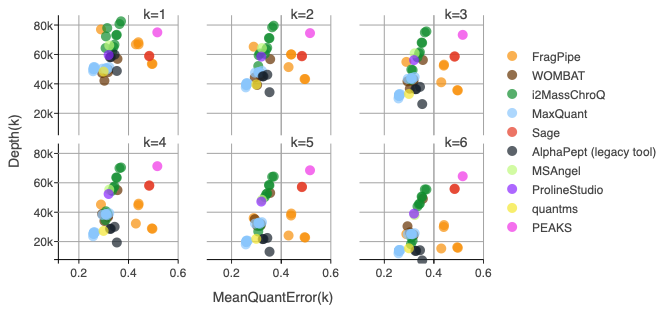

**Supplementary Figure 7:**  **Main figure of DDA Module, showing the MeanQuantError for k=1-6 minimum quantified ions.**

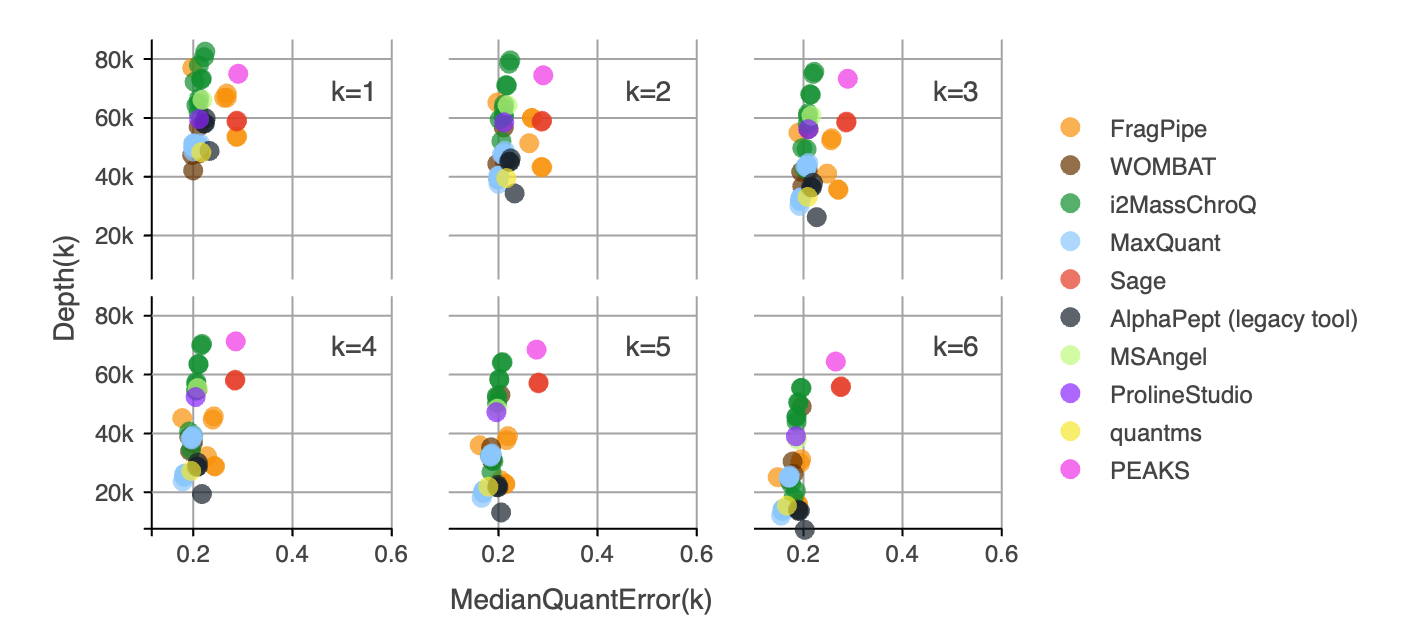

**Supplementary Figure 8:** **Main figure of DDA Module, showing the MedianQuantError for k=1-6 minimum quantified ions.**

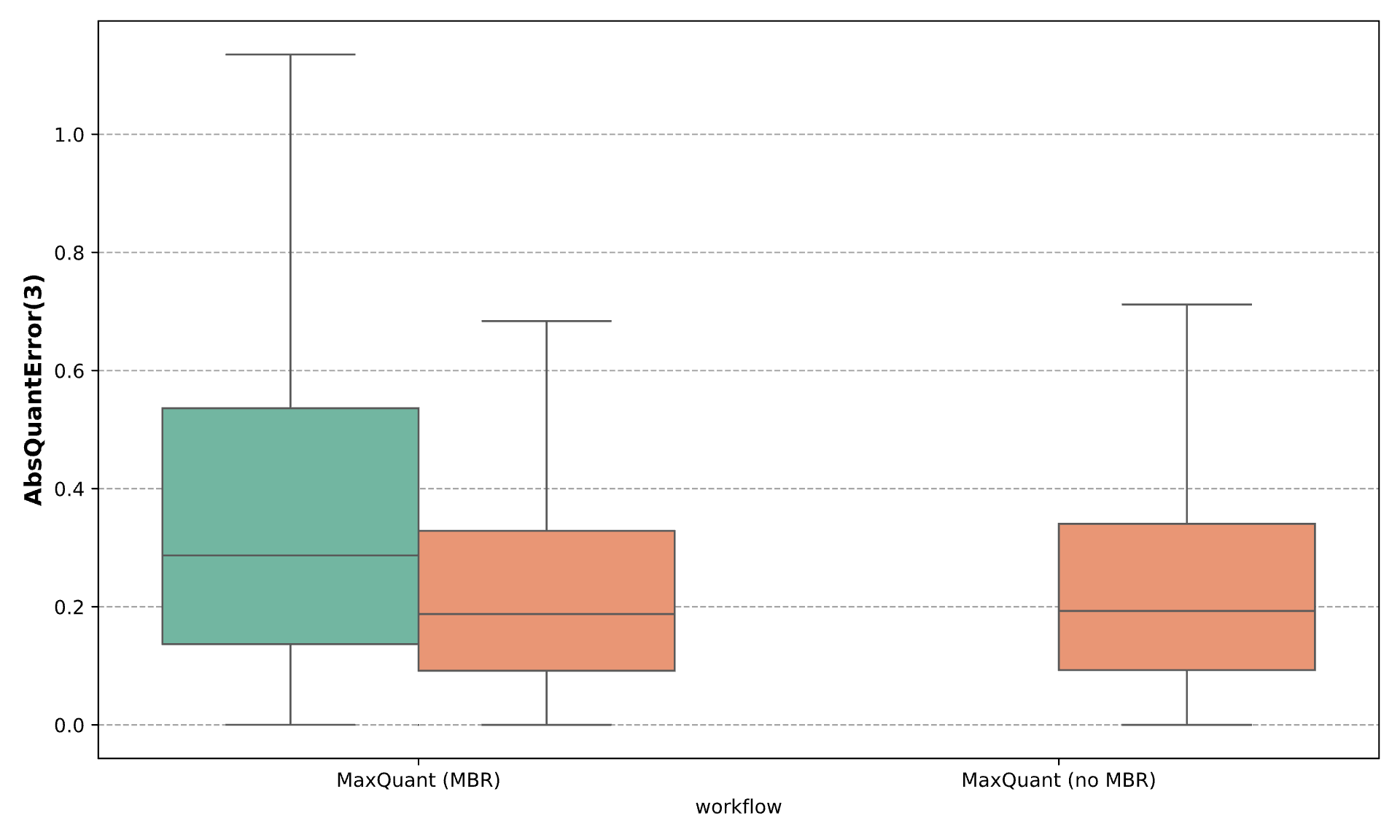

**Supplementary Figure 9:** **Impact of MBR on global precursor quantification error.** Boxplots of the AbsQuantError(3) of the precursor ions found in at least three replicates uniquely identified in the run with MBR enabled (green) or identified in both runs with MBR enabled (red, left) and without MBR enabled (red, right). “MaxQuant (MBR)”: MQ_B, “MaxQuant (no MBR)”: MQ_A.

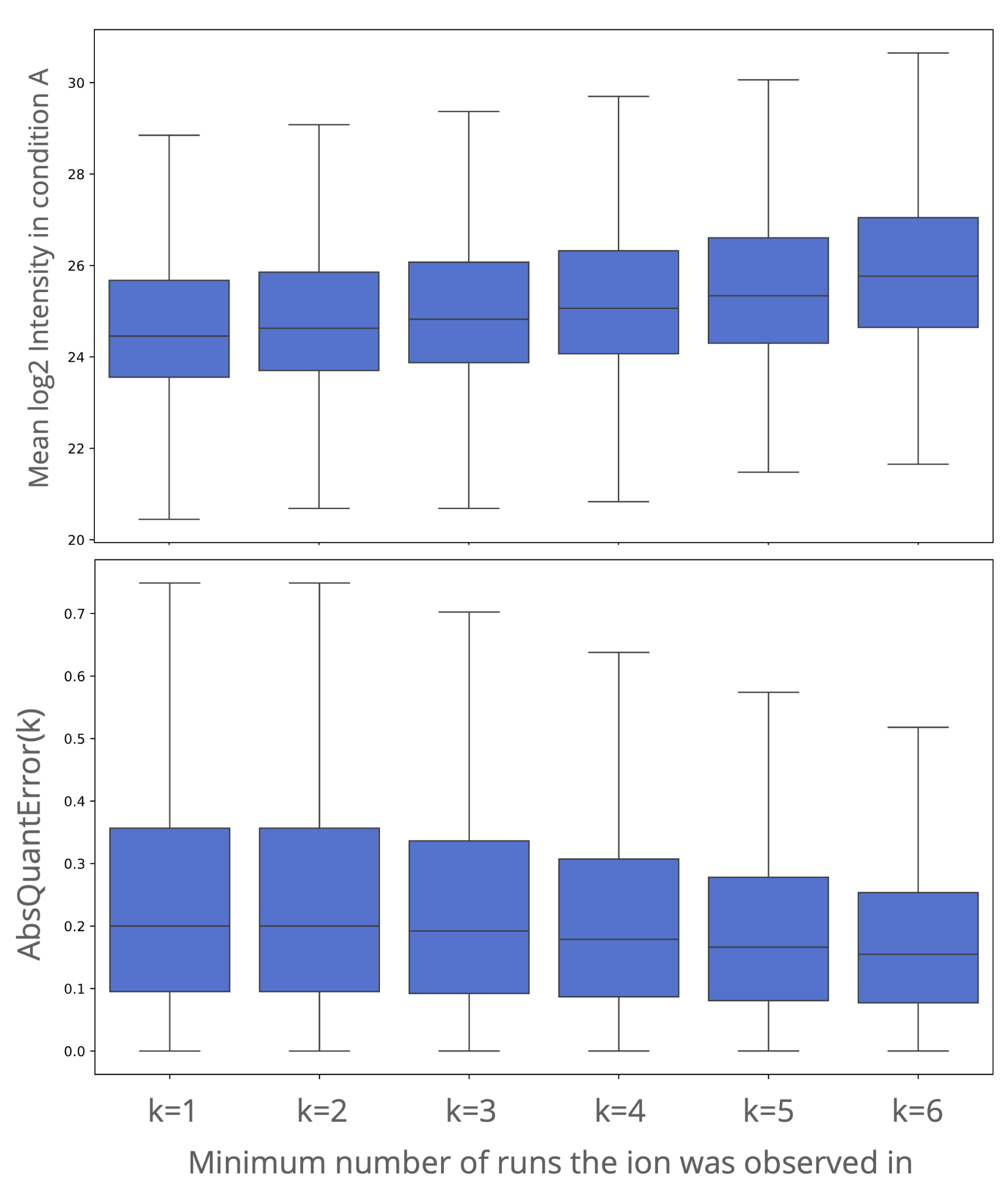

**Supplementary Figure 10:** **Precursor ions observed in most or all runs generally have higher intensity values and are better quantified.** Boxplot of the precursors log_2_-transformed mean intensity in condition A (top panel) and AbsQuantError (bottom panel) calculated for k = 1 to 6 (horizontal axis). Example data shown for condition A from the workflow MQ_C (MaxQuant run without MBR).

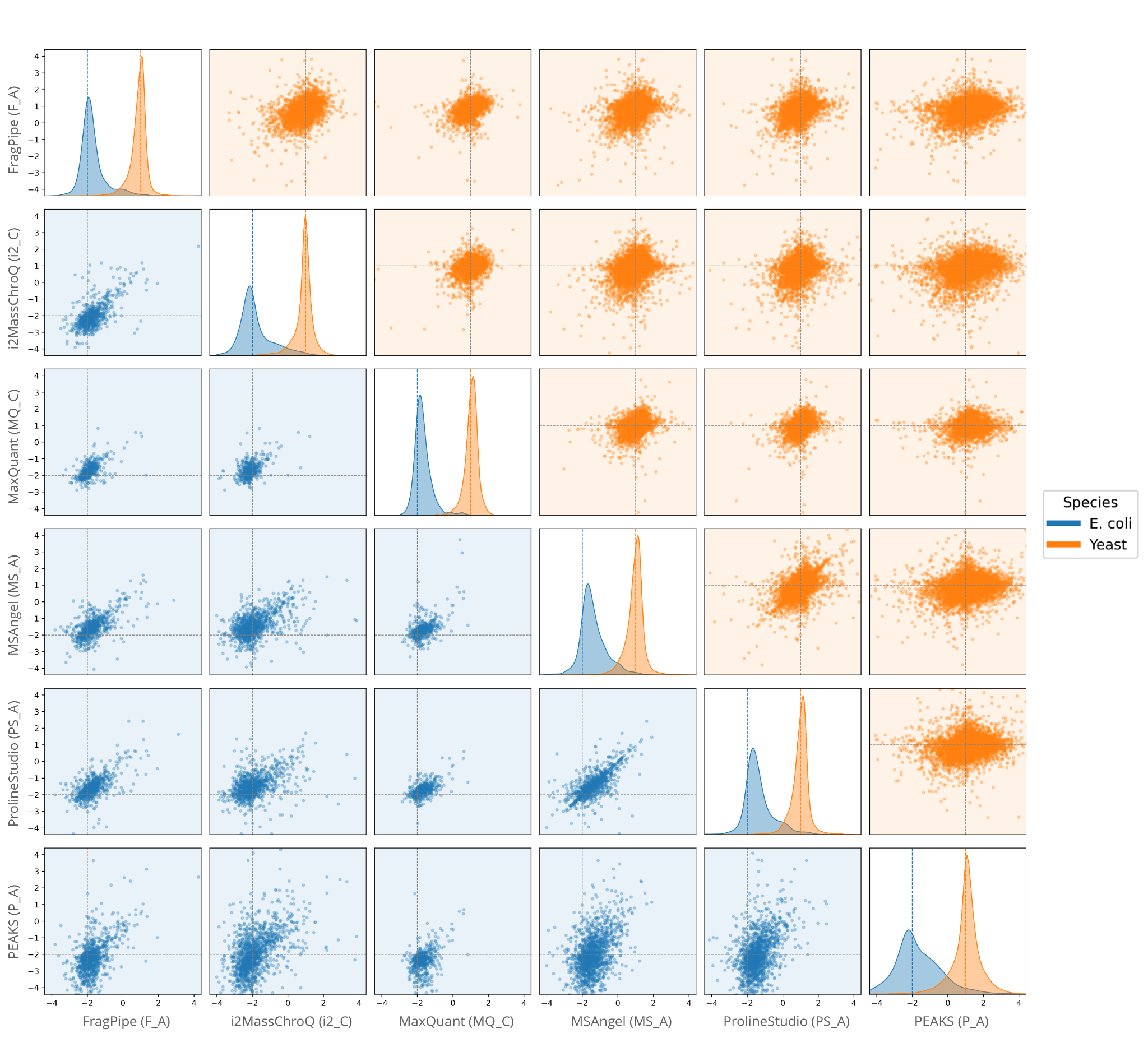

**Supplementary Figure 11:** **Pairwise comparison of precursor ion quantifications across software tools, with the QExactive DDA data.** Each dot represents the log2FC measured for a precursor ion between the two experimental conditions, as quantified by the indicated workflow. The density plots on the diagonal show the distribution of observed log2FC for each tool, overlaid by the expected log_2_-transformed (fold change (dotted line). The scatter plots display the pairwise correlation of fold change estimates for individual precursor ions between tools. Data are stratified by species, with *E. coli* shown in blue and yeast in orange.

**Supplementary Figure 12: Main figure of the DIA Astral Module, showing the MedianQuantError for k=1-6 minimum quantified ions.**

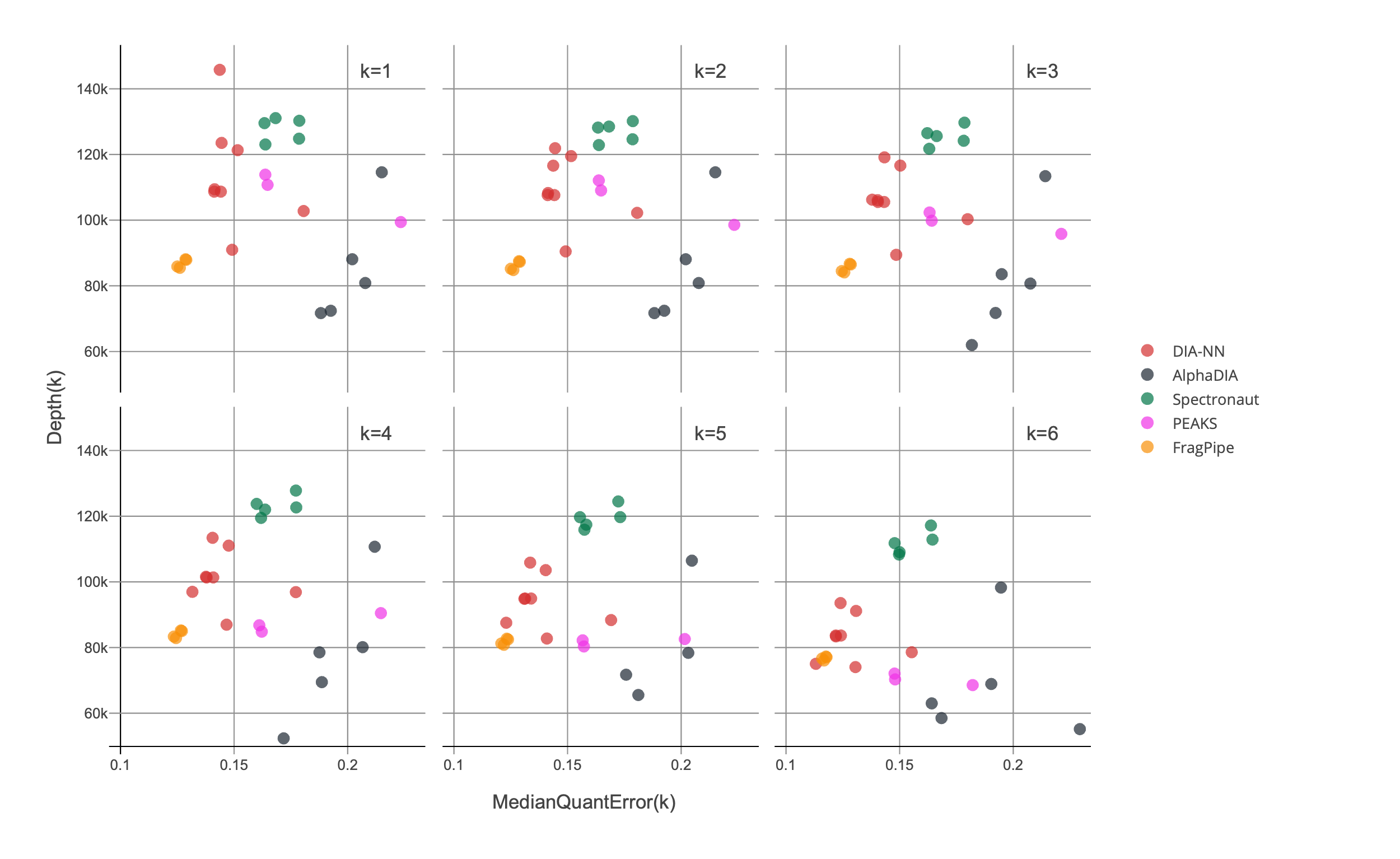

**Supplementary Figure 13: Main figure of the diaPASEF Module, showing the MedianQuantError for k=1-6 minimum quantified ions.**

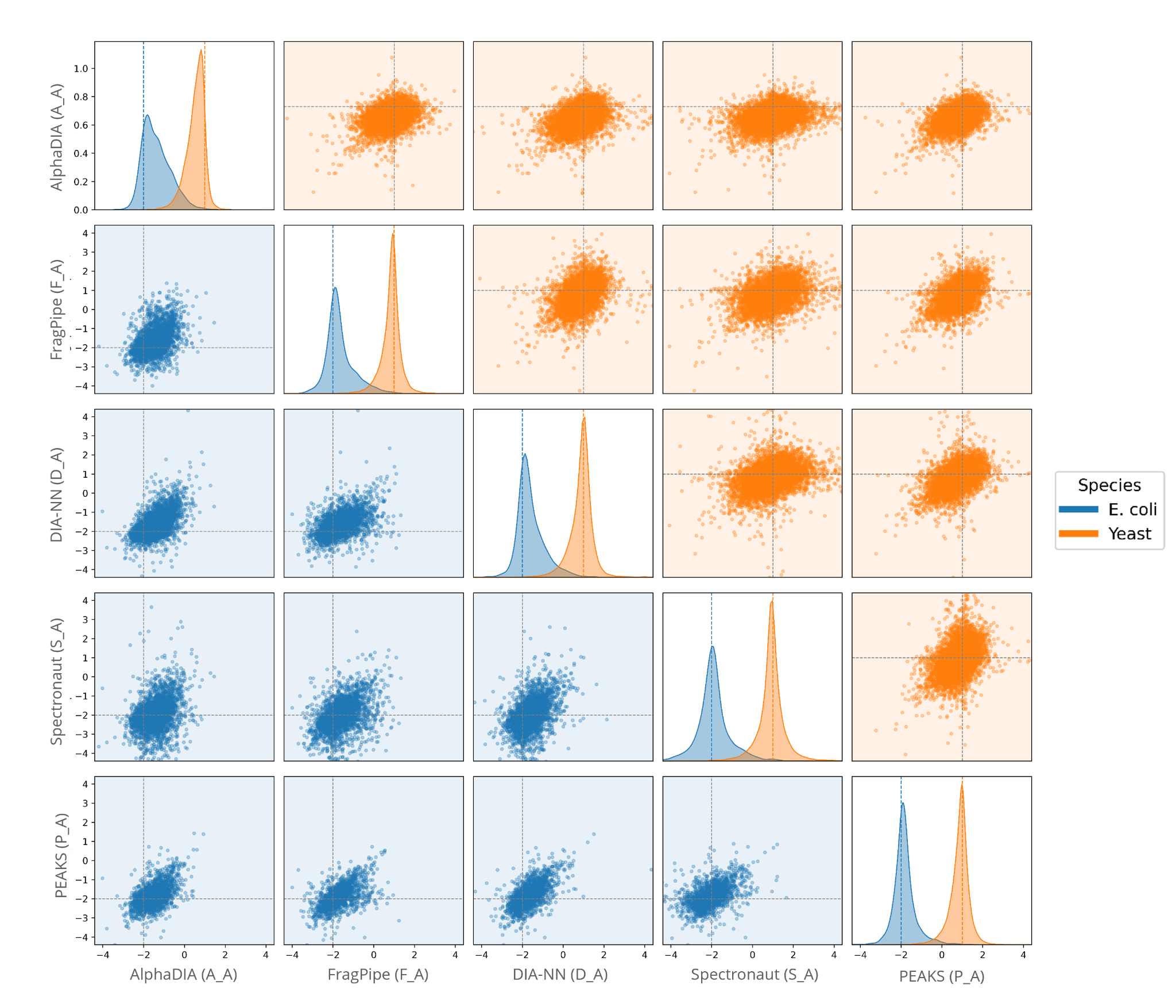

**Supplementary Figure 14:** **Pairwise comparison of precursor ion quantifications across software tools, with the Astral DIA data.** Each dot represents the log2FC measured for a precursor ion between the two experimental conditions, as quantified by the indicated workflow. The density plots on the diagonal show the distribution of observed log2FCs for each tool, overlaid by the expected log2FC (dotted line). The scatter plots display the pairwise correlation of fold change estimates for individual precursor ions between tools. Data are stratified by species, with *E. coli* shown in blue and yeast in orange. For improved visualization, the 0.5% most extreme outlying points across all comparisons were excluded from the plots, as their inclusion would otherwise excessively expand the axis ranges.
