## Supplementary Table 1 for "ProteoBench: the community-curated platform for comparing proteomics data analysis workflows"

**Supplementary Table 1:** Description of all the modules currently available (green background) or in development (blue background), and a selection of the most mature module proposals (orange background) in ProteoBench at time of submission. All associated descriptions and discussions are available here: github.com/orgs/Proteobench/discussions.

| **Module name** | **Description** | **Associated data** | **Level** |
| --- | --- | --- | --- |
| Quant LFQ DDA ion QExactive | Benchmark the ion-level quantification accuracy of LFQ workflows using a multi-species sample. | PXD028735: DDA data acquired on a Q Exactive HF-X Orbitrap (ThermoFisher Scientific)^1^. | Precursor ions |
| Quant LFQ DIA ion diaPASEF | Benchmark the ion-level quantification accuracy of LFQ workflows using a multi-species sample. | PXD062685: diaPASEF acquired on a timsTOF SCP (Bruker). | Precursor ions |
| Quant LFQ DDA peptidoform | Benchmark the peptidoform-level quantification accuracy of LFQ workflows using a multi-species sample. | PXD028735: DDA data acquired on a Q Exactive HF-X Orbitrap (ThermoFisher Scientific)^1^. | Peptidoforms |
| Quant LFQ DDA ion Astral | Benchmark the ion-level quantification accuracy of LFQ workflows using a multi-species sample. | DDA data acquired on an Orbitrap Astral (ThermoFisher Scientific). Full description and raw files available on the ProteoBench documentation website. | Precursor ions |
| Quant LFQ DIA ion Astral | Benchmark the ion-level quantification accuracy of LFQ workflows using a multi-species sample. | DIA data acquired on an Orbitrap Astral (ThermoFisher Scientific). Full description and raw files available on the ProteoBench documentation website. | Precursor ions |
| Single-cell label free DIA quantification | Benchmark the ion-level quantification accuracy of LFQ DIA workflows using a low-input multi-species sample. | In discussion: two proteome mix of HeLa and yeast acquired on an Orbitrap Astral (ThermoFisher Scientific)^2^. | Precursor ions |
| *De novo* DDA identification | Benchmark *de novo* peptide sequencing tools on data-dependent acquisition data. Evaluation is across different species and without any (FASTA) sequence information. | Curated balanced nine-species dataset designed for benchmarking of *de novo* identification algorithms^3^. All runs were acquired with DDA on Orbitrap instruments (Thermo Fisher Scientific). | Amino acids  Peptides |
| DDA protein identification and quantification with dynamic organellar maps | Benchmark identifications and LFQ workflows using subcellular co-fractionation (Dynamic Organellar Maps^4^). | PXD034971: DIA data acquired on an Exploris 480 (Thermo Fisher Scientific)^4^. | Protein groups |
| DDA identification - phosphopeptides | Benchmark phosphopeptide identification workflows and assess their identification and localization accuracy. | In discussion. | In discussion |
| DIA module on ZenoSWATH acquisition | Benchmark the ion-level quantification accuracy of LFQ workflows using a multi-species sample. | In discussion: data acquired on a ZenoSWATH instrument (SCIEX). | Precursor ions |
| LFQ in Human plasma | Benchmarking with a high dynamic range a multispecies dataset. | In discussion: data from Distler *et al*^5^. | Precursor ions |

Abbreviations: LFQ = label-free quantification; DDA = data-dependent acquisition; DIA = data-independent acquisition.
